# Forkhead transcription factor FKH-8 is a master regulator of primary cilia in *C. elegans*

**DOI:** 10.1101/2021.09.14.460205

**Authors:** Rebeca Brocal-Ruiz, Ainara Esteve-Serrano, Carlos Mora-Martinez, Juan Tena, Nuria Flames

**Affiliations:** Developmental Neurobiology Unit, Instituto de Biomedicina de Valencia IBV-CSIC, Valencia, 46010, Spain; Centro Andaluz de Biología del Desarrollo (CABD), Consejo Superior de Investigaciones Científicas/Universidad Pablo de Olavide, Seville, Spain

**Keywords:** cilium, transcriptional regulation, FKH, RFX, terminal differentiation, *C*. *elegans*

## Abstract

Cilia, either motile or non-motile (a.k.a primary or sensory), are complex evolutionary conserved eukaryotic structures composed of hundreds of proteins required for their assembly, structure and function that are collectively known as the ciliome. Ciliome mutations underlie a group of pleiotropic genetic diseases known as ciliopathies. Proper cilium function requires the tight coregulation of ciliome gene transcription, which is only fragmentarily understood. RFX transcription factors (TF) have an evolutionarily conserved role in the direct activation of ciliome genes both in motile and non-motile cilia cell types. In vertebrates, FoxJ1 and FoxN4 Forkhead (FKH) TFs work with RFX in the direct activation of ciliome genes, exclusively in motile cilia cell-types. No additional TFs have been described to act together with RFX in primary cilia cell-types in any organism. Here we describe FKH-8, a FKH TF, as master regulator of the primary ciliome in *Caenorhabditis elegans*. *fkh-8* is expressed in all ciliated neurons in C. *elegans*, binds the regulatory regions of ciliome genes, regulates ciliome gene expression, cilium morphology and a wide range of behaviours mediated by sensory cilia. Importantly, we find FKH-8 function can be replaced by mouse FOXJ1 and FOXN4 but not by members of other mouse FKH subfamilies. In conclusion, our results show that RFX and FKH TF families act as master regulators of ciliogenesis also in sensory ciliated cell types and suggest that this regulatory logic could be an ancient trait predating functional cilia sub-specialization.

## INTRODUCTION

Eukaryotic cilia are complex and highly organized organelles defined as specialized membrane protrusions formed from a stereotyped assembly of microtubules. Cilia are composed of hundreds of proteins, required for their assembly, structure and function, which are collectively known as the ciliome (**Figure 1A**). Cilia can be classified into motile or non-motile based on their function and structure: motile cilia are responsible for propelling cells or generating fluid flow while non-motile (a.k.a primary or sensory) cilia function as cellular antennae to sense extracellular stimuli (Choksi et al., 2014). Cilia appeared early in eukaryotic evolution and it is thought that ancient cilia displayed mixed motile and sensory functions (Mitchell, 2017). In multicellular invertebrates, primary and motor cilia are restricted to specific cell types. In contrast, in vertebrates, primary cilia are present almost in every cell, including neurons, while motile cilia are present only in specialized cell types.

**Figure 1.**
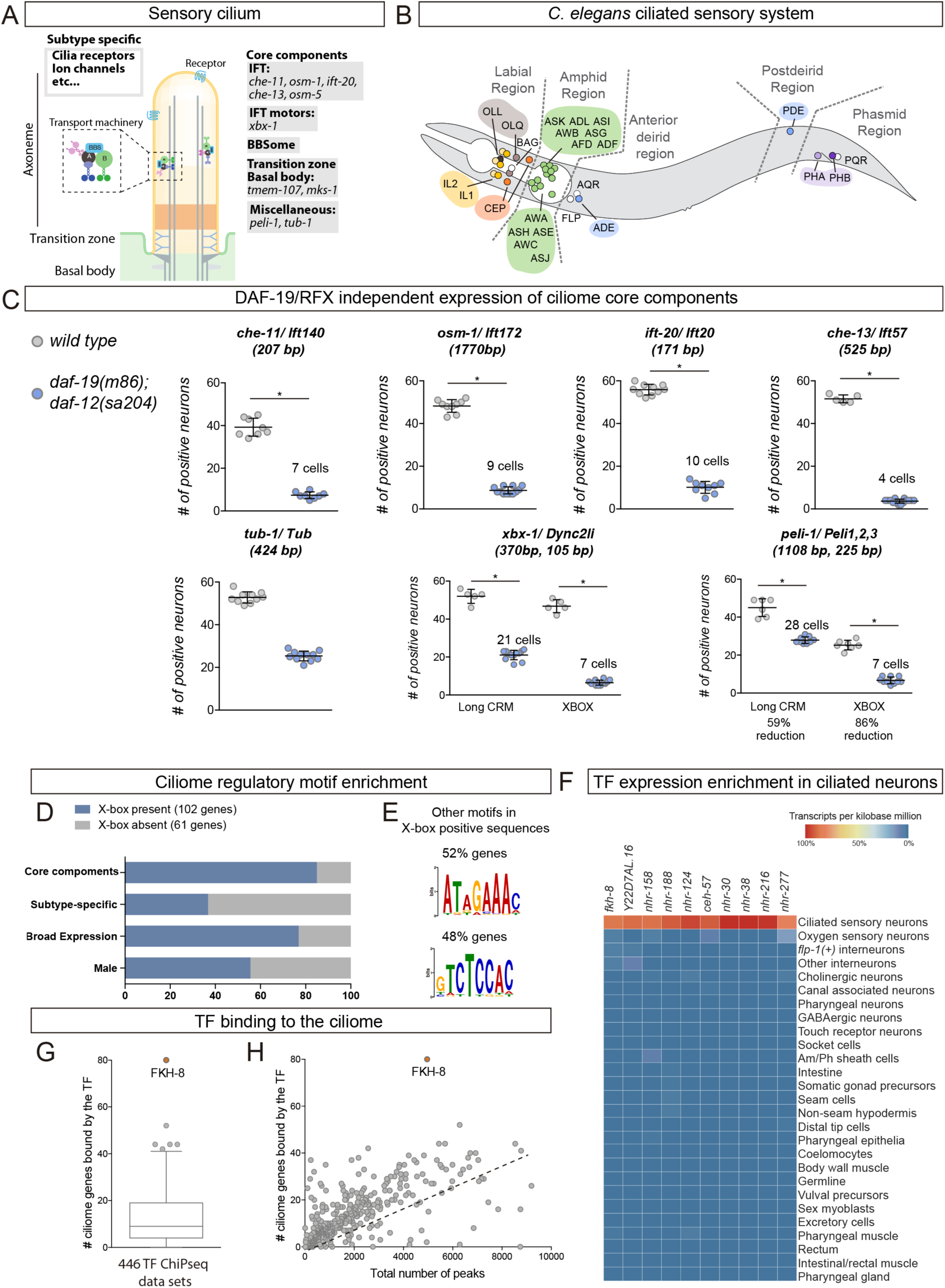
FKH-8 is a candidate regulator of ciliome gene expression in *C. elegans*. A) Schema for a sensory cilium. Cilia components (ciliome) can be divided into core and subtype specific categories. Genes whose reporters are analysed in panel C are indicated by their function. B) Left lateral view of *C. elegans* hermaphrodite ciliated system. Sixty ciliated neurons from 25 different classes are distributed in 5 distinct anatomical regions. C) Ciliome core components show persistent expression in double *daf-12(sa204), daf-19(m86)* null mutants. Each dot represents the total number of reporter-positive neurons in a single animal. Mean and standard deviation are represented. The mean number of remaining reporter-positive neurons in double *daf-12, daf-19* null mutants is indicated. See **Supplementary File 1** for raw data, *daf-12(sa204)* single mutant scorings and sample sizes and **Supplementary figure 2** for additional reporter scorings. D) DAF-19/RFX motifs (X-box) are enriched in regulatory sequences of core and broadly expressed ciliome genes. See **Supplementary figure 3** for additional enriched motifs. E) Motifs enriched in regulatory sequences of ciliome genes containing X-box sites, potential TF binding to these motifs is unknown. F) sc-RNA-seq data analysis (Cao et al., 2017) identifies 10 TFs specifically enriched in ciliated sensory neurons. These TFs belong to FKH, ZF, NHR and HD families. See **Supplementary figure 3** for detailed description of TF expression in each ciliated neuron type. G) ChIP-seq data analysis of 259 available TFs shows that FKH-8 ranks first in direct binding to genes of the ciliome list. See **Supplementary figure 3** for core ciliome or subtype specific binding. Correlation of total number of peaks and ciliome-list genes bound by TFs shows FKH-8 behaves as an outlier, demonstrating high binding to ciliome genes is not merely due to the high number of FKH-8 binding-events.

Most ciliome components are shared between motile and primary cilia and are referred as “core” ciliome (**Figure 1A**). In addition, motile cilia usually contain specialised axonemal dyneins and other motile-specific components while membrane of sensory cilia is decorated with receptors that trigger downstream signalling cascades when they are activated by small molecules, mechanical perturbations, or radiation.

The importance and wide range of cilia functions are underscored by the large number of congenital disorders caused by mutations in genes coding for ciliome components, which are collectively called ciliopathies (Andreu-Cervera et al., 2021; Horani and Ferkol, 2021; Lucas et al., 2020; Tobin and Beales, 2009). These disorders cause a broad spectrum of symptoms including retinal degeneration, polycystic kidney, deafness, polydactyly, brain and skeletal malformations, infertility, morbid obesity and mental retardation. Importantly, there are still many “orphan ciliopathies”, which correspond to congenital disorders classified as ciliopathies by phenotype but with yet unidentified causal mutations. Genetic variants lying in coding genes (including mutations in the ciliome) are easier to identify as causal mutations, however, most variants associated to human diseases lie in the non-coding genome. It is currently thought that some of these non-coding variants act as regulatory mutations affecting gene expression. Thus, regulatory mutations affecting ciliome gene expression might underlie many orphan ciliopathies, understanding the molecular mechanisms that ensure correct co-regulation of ciliome genes is then of utmost importance.

Little is known about the direct transcriptional co-regulation of ciliome gene expression (Choksi et al., 2014; Lewis and Stracker, 2020; Thomas et al., 2010). In 2000, pioneer work in *Caenorhabditis elegans* identified DAF-19, an RFX family transcription factor (TF), as a direct regulator of ciliome gene expression in the ciliated sensory neurons (Swoboda et al., 2000). This work was followed by numerous reports on the role of different members of the RFX TF family as direct ciliome regulators both in primary and motile cilia cell types in several animal models including *Drosophila melanogaster*, *Danio rerio*, *Xenopus laevis* and *Mus musculus* (Ashique et al., 2009; Bonnafe et al., 2004; Chung et al., 2012; Dubruille et al., 2002; Liu et al., 2007). FOXJ1, an ancient member of the Forkhead family, also acts as a direct activator of ciliome transcription in several vertebrates, but its role is limited to cell types containing motile cilia (Brody et al., 2000; Chen et al., 1998; Stubbs et al., 2008; Yu et al., 2008). Thus, currently additional TFs acting together with RFX TFs in the direct regulation of the ciliome in sensory cilia cell types are completely unknown in any organism.

Here, we take advantage of the amenability of *C. elegans* for genetic studies to understand the transcriptional regulatory logic of the non-motile primary cilliome. *C. elegans* contains sensory but not motile cilia. In hermaphrodites, sensory cilia are found in 25 out of the 118 neuronal types known as the ciliated sensory system (Scholey, 2007) (**Figure 1B**). We find that FKH-8, a FKH TF, is expressed in all ciliated sensory neurons in *C. elegans*, with an onset of expression concomitant to the start of ciliome gene expression. Chromatin immuno-precipitation and sequencing (ChIP-seq) data analysis shows that FKH-8 binds to a broad range of ciliome genes, at locations often near X-box sites, which are recognized by DAF-19/RFX. *fkh-8* mutants show decreased ciliome reporter gene expression, cilia morphology abnormalities and deficits in a wide range of behaviours mediated by sensory cilia. In addition, we find *fkh-8*/FKH and *daf-19*/RFX act synergistically in the regulation of ciliome genes. Finally, we show that mouse FoxJ1 and FoxN4, two ancient FKH TFs known to directly regulate ciliome expression in vertebrate motile-cilia cell types, rescue *fkh-8* mutant expression defects. This functional conservation is not observed with members of other FKH sub-families. Our results identify FKH-8 as the first TF acting together with RFX TFs in the direct regulation of the ciliome in sensory-ciliated cells and suggest that this function could be evolutionary conserved in vertebrates. Taken together, a global ciliome regulatory logic starts to emerge in which RFX and FKH TFs could act together in the direct regulation of ciliome gene expression both in cell types containing motile or primary cilia. Considering that ancestral eukaryotic cilium is proposed to combine motile and sensory functions, we speculate that RFX / FKH regulatory module might represent the ancestral state of eukaryotic ciliome gene regulation.

## RESULTS

### Persistent enhancer activity of ciliome genes in *daf-19*/RFX mutants

The activity of enhancers for ciliome genes bearing X-boxes is dramatically reduced in *daf-19(m86)* null mutants. However, for several ciliome fluorescent reporters, some residual activity has been anecdotally reported (Burghoorn et al., 2012; Chu et al., 2012; Efimenko et al., 2005; Haycraft et al., 2001; Stasio et al., 2018; Swoboda et al., 2000). As *daf-19* is the only RFX TF coded in the *C. elegans* genome, we reasoned that persistent ciliome enhancer activity in *daf-19(m86)* null mutants would underscore the presence of additional TF families working in concert with DAF-19. Based on previous data, we selected enhancers and built fluorescent reporters for ten phylogenetically conserved and broadly expressed core cilia components: five intraflagelar transport (IFT) genes (*che-11, osm-1, ift-20, che-13, osm-5*); the transition zone transmembrane genes *tmem-107* and *mks-1*; *a* Tub gene involved in receptor trafficking (*tub-1)*; the dynein-component *xbx-1* and the ubiquitin protein ligase *peli-1* (**Figure 1A**). Human orthologs for several of these genes are linked to ciliopathies (Horani and Ferkol, 2021; Mukhopadhyay et al., 2005; Thevenon et al., 2016). All fluorescent reporters contain at least one experimentally validated X-box (**Figure S1**). In *wild type* worms, all these reporters show broad activity in the ciliated system, with mean reporter expression in at least 30 ciliated neurons, except for *mks-1* and *osm-5* reporters that showed expression in less than 20 cells, suggesting other enhancers outside the analysed sequences might drive expression in additional ciliated neurons (**Figure 1C** and **Figure S2**). To avoid dauer entry of *daf-19(m86)* null animals, we analysed reporter expression in *daf-19(m86); daf-12 (sa204)* double mutants and added *daf-12 (sa204)* single mutant analysis as additional control (**Supplementary File 1**). As expected, *daf-19(m86); daf-12 (sa204)* double mutants show a dramatic decrease in the number of neurons positive for each reporter (**Figure 1C** and **Figure S2**). Importantly, all reporters except *tmem-107, mks-1* and *osm-5*, which correspond to the shortest constructs, show persistent expression in some neurons (**Figure 1C**, **Figure S2** and **Supplementary File 1**). We hypothesised that these short constructs might lack binding sites for additional TFs working with DAF-19. Indeed, we find that shorter versions of *xbx-1* and *peli-1* reporter constructs are more affected by *daf-19* mutation than corresponding longer constructs, consistent with shorter sequences lacking additional regulatory information (**Figure 1C**).

Unexpectedly, we find that *daf-12* itself has a small but significant effect on the expression of several reporters (namely, *che-13*, *ift-20*, *osm-1, mks-1* and *tmem-107*) (**Supplementary File 1**), suggesting a possible role for this nuclear hormone receptor (NHR) TF in ciliome expression. Altogether our data strongly suggests that additional TF or TFs act together with DAF-19 to directly activate core ciliome gene expression.

### Identification of FKH-8 as candidate regulator of ciliome gene expression

We reasoned that similar to *daf-19,* additional regulators of cilia gene expression could act broadly on many genes coding for ciliome components and in many different ciliated neuron types. Thus, to identify these putative candidates, we combined three strategies: *cis*-regulatory analysis of the ciliome genes, TF expression enrichment in the sensory ciliated system and TF binding to putative regulatory regions of the ciliome genes.

We built a manually curated list of 163 cilium effector genes (See materials and methods and **Supplementary File 2**). This list can be divided in four categories: 1) 73 “*core components*” present in all types of cilia and thus expressed by all ciliated neurons in *C. elegans*. Core components include IFT particles, kinesins, dyneins, BBSome complex, etc; 2) 68 “*Subtype specific*” genes, that code for channels or receptors located in cilia that are expressed in a neuron type specific manner, providing neuron type specific functions; 3) 13 “*Broad expression*” genes, specifically expressed within the ciliated system but not associated with a well-defined core cilia functions and 4) 9 “*Male*” genes that code for genes with male-specific cilia functions (**Supplementary File 2**).

D*e novo* motif enrichment analysis using the promoters of these ciliome genes identified previously known RFX consensus binding sites (X-box motif). In agreement with published results, X-box motifs are preferentially associated to “*Core”* and “*Broadly expressed”* ciliome genes (**Figure 1D**) (Burghoorn et al., 2012; Efimenko et al., 2005; Swoboda et al., 2000). An additional motif matching the pro-neural bHLH TF *lin-32*/Atoh1 is present in 28% of the genes, with no particular bias to any ciliome category (**Figure S3**). The pro-neural binding site might reflect the neuronal nature of this gene set, as in *C. elegans* cilia are only present in neurons. Motif enrichment analysis limited only to the 102 genes containing predicted X-box sites identified two additional motifs (**Figure 1E**). Neither of both motifs significantly match known Position Weight Matrices for TFs. In contrast to the X-box, which is highly specific, TF binding motifs (TFBM) for many TF families are often short and degenerate, thus they appear widely in the genome and provide low information content. This feature might explain the failure to find enriched motifs for additional TFs in ciliome gene regulatory regions.

As an alternative to motif enrichment analysis, we turned to TF expression enrichment. We hypothesized that TFs acting broadly on sensory cilia gene expression could show enriched expression in the sensory ciliated neurons. Using available single cell RNA expression data (sc-RNAseq) from the second larval stage (Cao et al., 2017), we retrieved the expression pattern of 861 *C. elegans* transcription factors (Narasimhan et al., 2015). Ten transcription factors are specifically enriched within the ciliated sensory neurons compared to other neuron types or non-neuronal tissues (**Figure 1F**). Using an independent set of sc-RNAseq data from young adult (Taylor et al., 2021), only FKH-8 expression is detected in all 25 different types of sensory ciliated neurons (**Figure S3**), suggesting it could be a good candidate to work together with DAF-19.

Finally, we interrogated 446 published ChIP-seq datasets, corresponding to 259 different TFs (including FKH-8 but not DAF-19), for nearby binding to the ciliome gene list (**Supplementary File 2**). We find FKH-8 behaves very differently from the rest of TFs with at least one FKH-8 binding peak associated to 49% of the genes on the ciliome gene list (**Figure 1G-H**). FKH-8 binds both core components and subtype specific ciliome genes (**Figure S3**), although, similar to X-box motifs, FKH-8 binding is significantly more prevalent for core ciliome genes (75% compared to 22% binding to subtype genes). Thus, both sc-RNAseq and ChIPseq data analysis pinpoint FKH-8 as a good candidate TF to directly control ciliome gene expression.

### FKH-8 is expressed in all ciliated sensory neurons

FKH-8::GFP fosmid expression at young adult stage is detected in all ciliated sensory neurons, as assessed by co-localization with the *ift-20* core ciliome reporter (**Figure 2A, B and Supplementary File 1**). Only three non-ciliated neurons, PVD, VC4 and VC5 show FKH-8 expression, while no expression is detected in non-neuronal tissues. *C. elegans* male nervous system contains additional ciliated sensory neurons, mostly in the tail, which also express FKH-8 (**Figure 2B**). During embryonic development, there is a similar overlap between FKH-8 and *ift-20* reporters (**Figure 2C and S4**). Correlation between *fkh-8*, *ift-20* and *daf-19* expression during development is also observed using Uniform Manifold Approximation and Projection (UMAP) representation of embryonic sc- RNA-seq data (Packer et al., 2019) (**Figure 2D and S4**). In addition, there is a high gene expression correlation for the 73 core ciliome genes and *daf-19* or *fkh-8* expression but not with other TFs (**Figure S4**). Thus, our analysis shows that FKH-8 is expressed almost exclusively in the whole ciliated sensory system and its developmental expression correlates with core ciliome gene expression.

**Figure 2.**
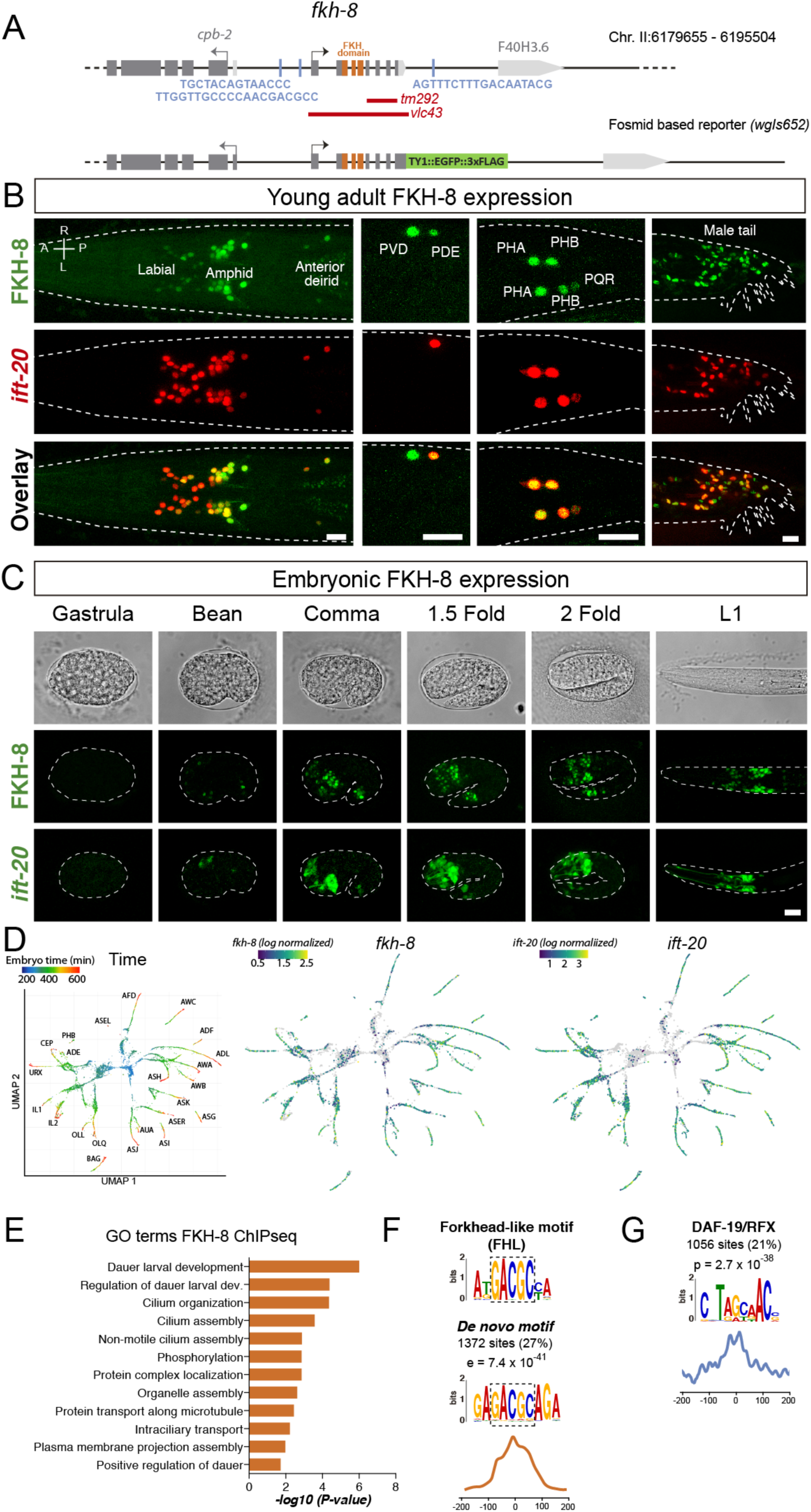
FKH-8 expression correlates with ciliated neuron differentiation. A) *fkh-8* locus (top) and fosmid based *fkh-8* reporter (bottom). Grey boxes represent exons and orange boxes correspond to exons coding for the FKH DNA binding domain. Putative *daf-19*/RFX binding sites are depicted with a blue bar. Red lines indicate extension for the corresponding deletion alleles. B) Dorso-ventral views of young adult animals expressing both the fosmid-based FKH-8::GFP reporter (in green) and an integrated reporter for the panciliary marker *ift-20* (in red). A: anterior, P: posterior, R: right, L: left. Scale bar = 10 µm. C) Embryonic developmental expression of FKH-8::GFP matches in time and space panciliary reporter *ift-20::gfp* expression. Scale bar = 10 µm. See **Supplementary figure 4** for embryonic co-localization between FKH-8::GFP and *ift-20::tagrfp* reporter. D) Embryonic sc-RNA-data (Packer et al., 2019) from *C. elegans* ciliated neurons and their progenitor cells. Pseudo-time (left pannel) shows the maturation trajectory of ciliated neurons that coincides with increasing *fkh-8* (centre) and *ift-20* (right) expression. See **Supplementary figure 4** for *daf-19* expression analysis and quantification of gene expression correlation. E) Genes bound by FKH-8 enrich Gene Ontology terms related to cilia regulated processes and/or functions. Data correspond to adjusted p-value. F) *De novo* motif analysis of FKH-8 ChIP-seq data identifies a motif present in 27% of the peaks, enriched at central peak positions, that matches an FHL-like motif. DAF-19/RFX binding motifs (PWM M1534_1.02) are present in 21% of the FKH-8 bound regions and are enriched at central positions. See **Supplementary figure 4** for similar analysis on additional FKH ChIP-seq data sets.

### FKH-8 genomic binding is enriched in ciliome genes

Next, we extended FKH-8 ChIPseq data analysis to the whole genome. FKH-8 binds a total of 5035 genomic regions assigned to 3,987 genes. Most peaks are associated to promoter regions (58,65%). Gene ontology analysis of FKH-8 bound genes shows enrichment for cilia functions or dauer regulation (which is also dependent on cilia integrity) (**Figure 2E**).

DNA consensus sequences bound by FKH-8 have not been experimentally determined. FKH TF family binds the canonical consensus RYMAAYA (Pierrou et al., 1994) and an alternative motif, termed FKH-like (FHL), characterized by a GACGC core sequence (Nakagawa et al., 2013). *De novo* motif enrichment analysis of FKH-8 ChIP-seq peaks does not show any match for FKH canonical binding site but identifies a motif that highly resembles the FHL motif (**Figure 2F**). This motif, present in 27% of the peaks, is enriched at central positions suggesting it could act as FKH-8 primary binding motif (**Figure 2F**).

We noticed that eight out of the twelve functional X-boxes present in the core ciliome reporters analysed in Figure 1C overlap with FKH-8 ChIP-seq peaks (**Figure S1**). Thus, we next looked for DAF-19 biding motif enrichment in FKH-8 bound regions. 21% of FKH-8 peaks contain at least one match for the DAF-19 position weight matrix (**Figure 2G**). Importantly, predicted X-boxes are preferentially found also at central locations, suggesting they could be in close proximity to FKH-8 bound sites (**Figure 2G**). DAF-19 binding sites are less significantly or not significantly enriched in ChIP-seq datasets for other FKH TFs (**Figure S4**), suggesting specific co-binding of DAF-19 and FKH-8.

### *fkh-8* mutants show defects in ciliome reporter gene expression

The only available *fkh-8* mutation, *tm292*, is a deletion downstream the FKH DNA binding domain, suggesting it could act as a hypomorphic allele (**Figure 2A**). Thus, we built *fkh-8(vlc43),* a null deletion allele that removes the whole *fkh-8* locus (**Figure 2A**). We selected eight reporters for six genes that code for core cilia components and that overlap with FKH-8 ChIP-seq peaks (**Figure S1**) and analysed their expression both in *fkh-8(tm292)* and *fkh-8(vlc43)* mutants.

Both *fkh-8* mutant alleles show significant expression defects in all reporters except for *tub-1*/Tub and the long *peli-1/*Peli1,2,3 reporter. Phenotypes are often more penetrant in *fkh-8(vlc43)* null allele than in the *fkh-8(tm292)* supporting the hypomorphic nature of *tm292* allele (**Figure 3A and Supplementary File 1**). The expression of each reporter is affected in specific subpopulations of ciliated neurons with some residual reporter expression found in all cases (**Figure 3A**). Lack of fluorescence reporter expression in *fkh-8* mutants reflects enhancer activity defects and not the absence of the ciliated neurons *per se*, as in *fkh-8* mutants *tub-1*/Tub and the long *peli-1/*Peli1,2,3 reporters are expressed in 53 and 46 ciliated neurons respectively (**Supplementary File 1**), similar to *wild type* expression levels.

**Figure 3.**
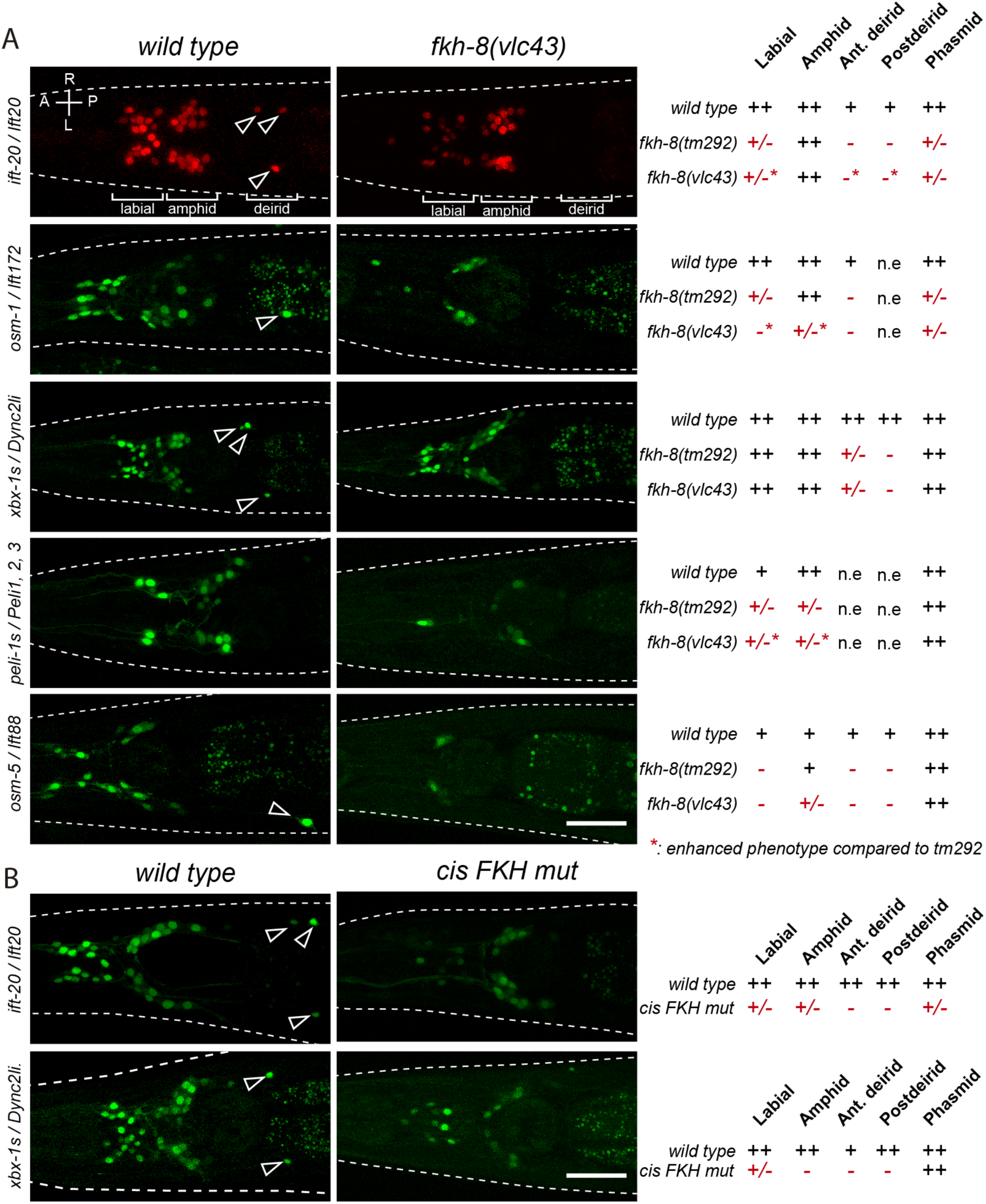
FKH-8 TF and FKH-binding sites are required for correct core ciliome gene reporter expression. A) Dorso-ventral images from young adult heads expressing different core ciliome gene reporters in *wild type* and *fkh-8(vlc43)* null mutant animals. Significant expression defects in 5 distinct anatomical regions for both *tm292* and *vlc43* alleles are summarized in the right panel. *Wild type* reporter expression “++” indicates more than 50% of ciliated neurons in the region express the reporter, whereas “+” indicates expression in less than 50% of the neurons. n.e: not expressed. Statistically significant expression defects appear in red. Expression defect below 50% of *wild type* values are indicated as “+/-” whereas losses higher than 50% are depicted as “-”. Enhanced phenotype in *vlc43* mutants compared to *tm292* is marked with an asterisk. A: anterior, P: posterior, R: right, L: left. Scale bar = 25 µm. See **Supplementary File 1** for raw scoring data. *Cis*-regulatory mutation of putative FKH sites greatly reduces ciliome gene reporter expression. Representative dorso-ventral images from young adult heads expressing *wild type* or FKH-site-mutated reporters for core ciliome genes *xbx-1* and *ift-20*. Expression defects are summarized on the right as in A). A: anterior, P: posterior, R: right, L: left. Scale bar = 25 µm. See **Supplementary Figure 5** and **Supplementary File 2** for *cis*-mutation details.

*fkh-8(vlc43)* animals show missing *ift-20* expression in ten neurons including the four pairs of dopaminergic ciliated mechanosensory neurons (CEPV, CEPD, ADE and PDE). Expression in *fkh-8(vlc43)* animals of *fkh-8* cDNA under the control of a *dat-1* dopaminergic specific promoter, which is unaffected in *fkh-8* mutants, is able to rescue *ift-20* reporter expression, consistent with a cell autonomously role of *fkh-8* in the regulation of ciliome gene expression (**Supplementary File 1,** see also **Figure 7**).

Next, we complemented the TF mutant analysis with *cis*-regulatory mutant analysis. We focused on *ift-20* and short *xbx-1* reporters which both overlap with FKH-8 Chip-seq peaks (**Figure S5**). Three independent transgenic lines with point mutations for FKH binding sites show broad expression defects both for *ift-20* and *xbx-1* reporters (**Figure 3B and Supplementary File 1**). *cis*-mutation expression defects are stronger than the ones observed for *fkh-8* mutant alleles suggesting either other FKH factors can compensate the lack of *fkh-8* or that *cis*-mutations could affect the binding of other TFs in addition to FKH-8.

In summary, our *cis* regulatory and *fkh-8* mutant analyses unravel a cell autonomous role for FKH-8 in the regulation of ciliome gene expression.

### FKH-8 is required for correct cilia morphology

Mutations in several ciliome core component, including *osm-5* and *xbx-1*, whose reporters are affected in *fkh-8* mutants, show cilium morphology defects (Blacque et al., 2004; Mukhopadhyay et al., 2007; Perkins et al., 1986; Starich et al., 1995). One of the most commonly used methods to assess gross cilium integrity is lipophilic dye staining (like DiD), which in *wild type* animals labels a subpopulation of amphid and phasmid neurons (Starich et al., 1995). Unexpectedly, *fkh-8(vlc43)* animals show similar DiD staining than *wild type* (**Supplementary File 1**).

Next, we directly analysed cilium morphology labelling specific subpopulations of ciliated neurons (**Figure 4**). Cilium length in CEP and AWB neurons is significantly reduced in *fkh-8(vlc43)* mutants compared to controls, while ADF neuron cilium length is significantly increased (**Figure 4**). In addition, *fkh-8* mutants display arborization defects in AWA cilia, while AWC cilia showed no major defects (**Figure 4**).

**Figure 4.**
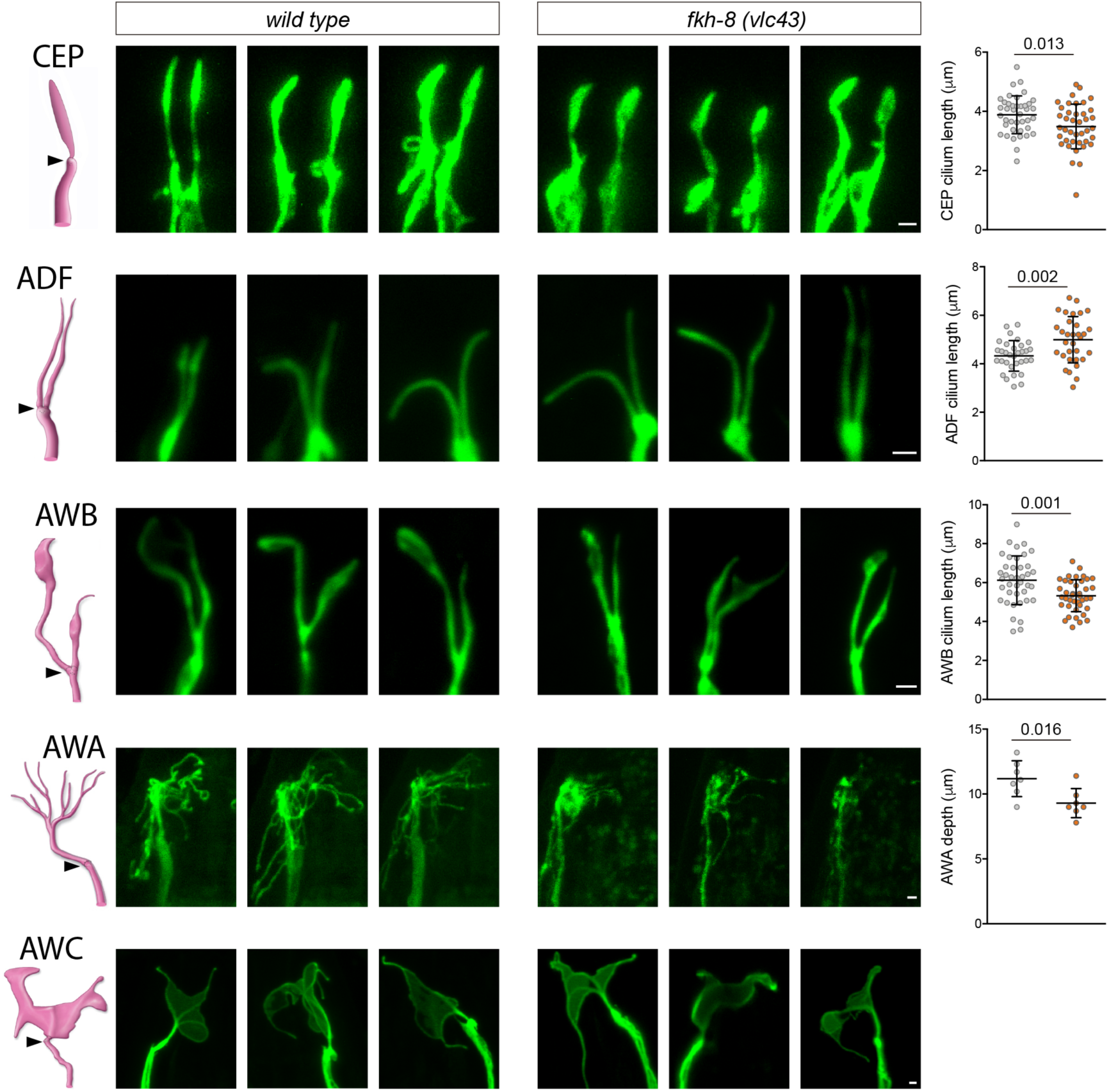
*fkh-8(vlc43)* null mutants display morphological defects in cilia. Integrated reporters unaffected in *fkh-8* mutant are used to label the cilia of several distinct subpopulations of ciliated neurons. CEP: *otIs259(dat-1::gfp)*; ADF: *zdIs13(tph-1::gfp)*; AWB: *kyIs104(str-1::gfp)*; AWA: *pkIs583(gpa-6::gfp)*; AWC: *kyIs140(str-2::gfp)*. Panels show representative images from 3 animals in *wild type* and *fkh-8(vlc43)* mutant backgrounds. Cilium length of CEP and AWB neurons is significantly reduced in the absence of FKH-8 whereas ADF cilia length is increased. Depth of AWA cilium arborization is significantly reduced in *fkh-8(vlc43)* null mutants. No major defects are observed in AWC cilia when comparing both genetic backgrounds. Each dot in the graphs represents measures for a single cilium. Mean and standard deviation are represented.

Thus, FKH-8 is necessary to regulate correct cilium length and morphology in diverse types of ciliated neurons.

### *fkh-8* mutants display defects in a wide range of cilia mediated behaviours

In *C. elegans* cilia are necessary to mediate sensory functions (Bargmann, 1993); thus, we interrogated *fkh-8* mutants with a battery of sensory paradigms. *fkh-8* mutants respond similarly to *wild type* animals to body gentle touch stimuli, which are mediated by not ciliated neurons (Chalfie and Sulston, 1981) (**Figure S6**), discarding general response or locomotory defects in *fkh-8* mutants. Response to posterior harsh touch, which is redundantly mediated by ciliated PDE and non-ciliated PVD neurons (Li et al., 2011) is also unaffected in *fkh-8(tm292)* and *fkh-8(vlc43)* animals, suggesting FKH-8 is not required to mediate this mechanosensory behaviour (**Figure S6**).

We tested two additional mechanosensory behaviours mediated only by ciliated sensory neurons: basal slowing response, mediated by dopaminergic ciliated neurons (Sawin et al., 2000) and nose touch, mediated by ASH, FLP and OLQ ciliated neurons (Kaplan and Horvitz, 1993). No defects on basal slowing response are found in *fkh-8(vlc43)* null mutants, while both *fkh-8* alleles are defective for nose touch responses (**Figure 5A, B)**. *vlc43* null allele shows stronger defects than *tm292* allele, supporting the hypomorphic nature of *tm292* allele (**Figure 5A)**.

**Figure 5.**
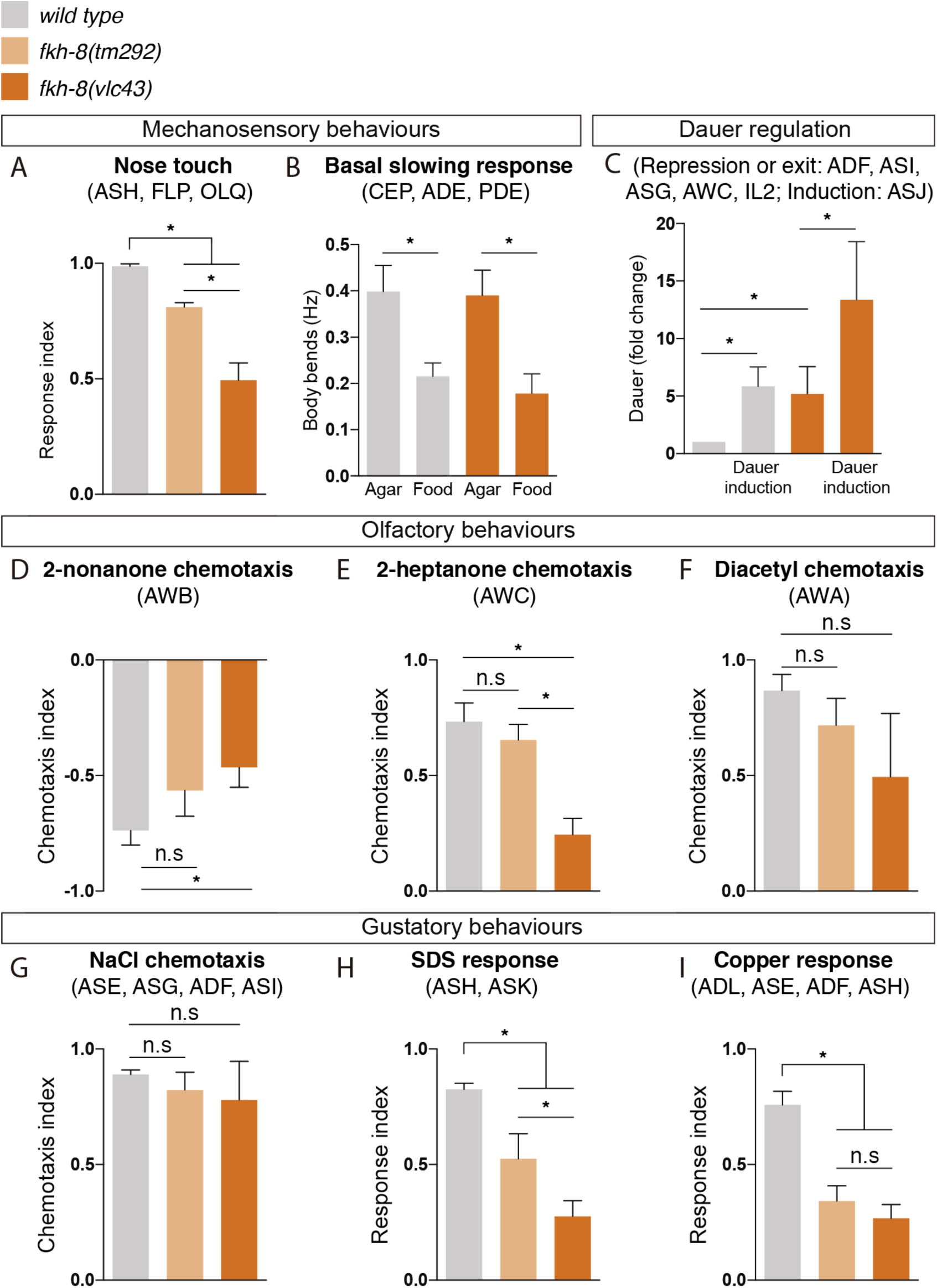
FKH-8 is required for the correct display of several sensory mediated behaviours. A) Mutations in *fkh-8* significantly impair appropriate backward response to nose touch, revealing functionality defects for the ASH, FLP and/or OLQ ciliated neurons. This phenotype is stronger in *fkh-8(vlc43)* null mutants than in the hypomorphic *tm292* allele. B) Decrease in locomotory rate upon re-entering a bacterial lawn is unaffected in *fkh-8* mutants. C) *fkh-8* null mutants significantly fail to prevent *dauer* entry. Pheromones induce *dauer* in *fkh-8* mutants, albeit less efficiently than in controls. D to F) Lack of *fkh-8* significantly impairs olfaction-mediated behaviours towards compounds sensed by ciliated AWB and AWC neurons. Diacetyl response, mediated by AWA, is affected but not to a significant level due to high variability in the response. G to I) Attractive chemotaxis towards NaCl is unaffected in *fkh-8* mutant animals. Avoidance behaviour towards toxic SDS and copper anions is significantly impaired. Mean and standard deviation are represented in all graphs. See **Supplementary figure 6** for quantification of non-cilia mediated behaviours and **Supplementary file 3** for raw data and samples’ sizes.

*fkh-8(vlc43)* animals are slightly but significantly dauer constitutive at 27°C compared to N2 controls (**Figure 5C**), indicating defects in preventing dauer entry, which is mediated by ADF, ASI and ASG ciliated neurons (Bargmann and Horvitz, 1991). Moreover, exposure to pheromones induces dauer entry in *fkh-8(vlc43)* animals less efficiently than in *wild type* animals [6 fold induction in *wild type* versus 3 fold induction in *fkh-8(vlc43)* animals] (**Figure 5C**), suggesting FKH-8 is also required for correct dauer entry, which is mediated by ASJ ciliated neuron (Bargmann and Horvitz, 1991).

*fkh-8(vlc43)* null mutants, but not *tm292* allele, show significant odor avoidance defects to 2-nonanone (AWB mediated) and defective odor attraction to 2- heptanone (mediated by AWC) (**Figure 5D, E**) (Bargmann et al., 1993; Sengupta et al., 1996; Troemel et al., 1997). Diacetyl attraction, which is mediated by AWA, is also decreased in *fkh-8(vlc43)* animals, although not significantly, due to high standard deviation values (**Figure 5F**).

Finally, we tested gustatory responses to NaCl, Sodium Dodecyl Sulfate (SDS) and copper. *fkh-8* mutants are attracted to NaCl similar to N2 controls, a response that is mediated mainly by ASE ciliated neurons (Bargmann and Horvitz, 1991) (**Figure 5G**). In contrast, avoidance response to SDS, mediated by ASH and ASK ciliated neurons (Hilliard et al., 2002) and avoidance to copper, mediated by ASH, ASE, ADF and ADL ciliated neurons (Guo et al., 2015; Sambongi et al., 1999), were significantly reduced both in *fkh-8(vlc43)* and *fkh-8(tm292)* animals (**Figure 5H, I**).

In summary, our battery of behavioural assays reveals FKH-8 is necessary for the correct response to a wide range of sensory stimuli (mechanical, gustatory or olfactory) that are mediated by different types of ciliated neurons (ADF, ADL, ASE, ASG, ASH, ASI, ASJ, ASK, AWB, AWC, FLP and OLQ). Specific behaviours, such as attraction to NaCl or basal slowing response are sustained in *fkh-8* mutants, suggesting retained sensory functions for particular neuron types, even with gene expression or morphological cilia defects (such as CEPs).

### FKH-8 and DAF-19/RFX act synergistically

FKH-8 binds five different locations in the *daf-19* locus (**Figure 6A**) while *fkh-8* locus contains 3 putative X-box sites (**Figure 2A**), suggesting they could cross-regulate each other’s expression. Transcription of *daf-19* locus generates different isoforms that share the carboxyl terminal (Ct) domain and the RFX DNA binding domain but differ in the amino-terminal region (**Figure 6A**). Some of these isoforms are expressed in a mutually exclusive manner: *daf-19d* is specifically expressed in ciliated neurons while *daf-19a/b* isoforms are expressed in the rest of the nervous system but not in ciliated neurons (Senti and Swoboda, 2008). Accordingly, a fosmid based Ct-tagged DAF-19 reporter that labels all isoforms is broadly expressed in neurons (**Figure S7**). We did not find any obvious DAF-19::GFP expression defects in *fkh-8(vlc43)* mutants (**Figure S7**). Co-localization of DAF-19::GFP with *dat-1::mcherry* dopaminergic reporter expression or DiD lipophilic staining also reveals similar expression in *wild type* and *fkh-8(vlc43)* mutants in the dopaminergic or amphid ciliated neurons (**Figure S7**). Thus, our data suggest that, despite its extensive binding to *daf-19* locus, FKH-8 does not seem to be required for *daf-19* expression, at least in the subpopulation of ciliated neurons directly assayed.

**Figure 6.**
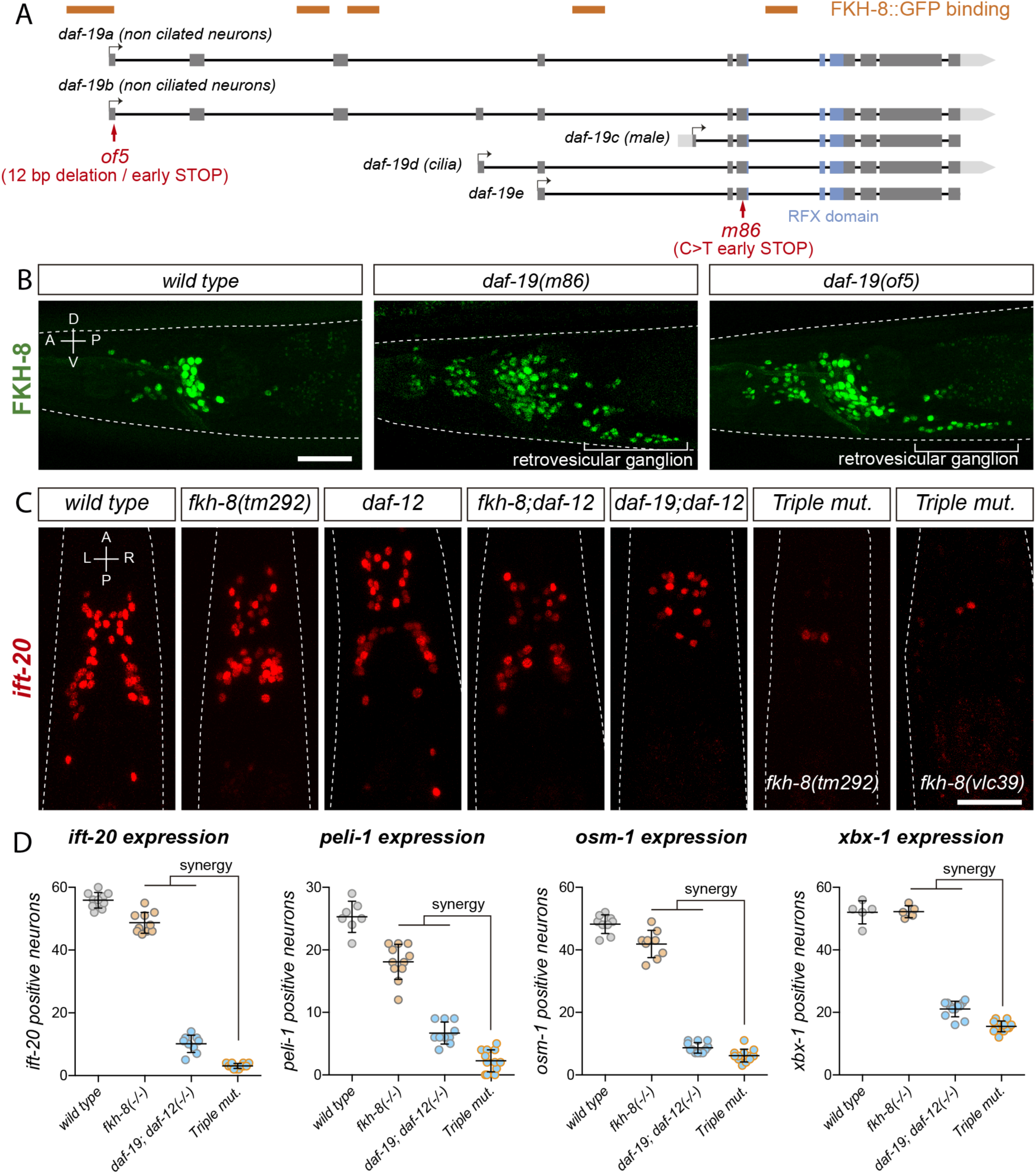
FKH-8 and DAF-19 exhibit crosstalk and synergistic effects in the transcriptional regulation of the ciliome. A) *daf-19* locus codes for five different *daf-19* isoforms. Grey boxes represent exons whereas blue boxes correspond to exons coding for the RFX DNA binding domain. FKH-8 binding events are depicted as orange bars. Red arrows locate mutations of the corresponding alleles. B) Lateral views from young adult hermaphrodite heads expressing the *fkh-8* fosmid-based reporter *wgIs652*. Lack of all *daf-19* isoforms (*m86* allele) derepresses *fkh-8* in non-ciliated neurons. This phenotype is mimicked by the specific absence of long *daf-19a/b* isoforms (*of5* allele). Scale bar = 25 µm. C) Dorso-ventral images from young adult hermaphrodites showing core ciliome *ift-20* reporter expression in several genetic backgrounds. Scale bar = 25 µm. Mean number of reporter-expressing neurons is significantly lower than expected in *fkh-8(tm292)*, *daf-19(m86)*, *daf-12(sa204)* triple mutants for the core ciliome features *ift-20*, *peli-1*, *osm-1* and *xbx-1*. Each dot represents the number of reporter-expressing neurons scored in a single animal. Mean and standard deviation are represented. See **Supplementary File 1** for raw data and statistics for all analysed genetic backgrounds.

Next, we assessed FKH-8::GFP fosmid expression in *daf-19(m86); daf-12(sa204)* double null mutants. In contrast to the pan-ciliated neuron expression pattern seen in *wild type,* FKH-8::GFP is expressed pan-neuronally in *daf-19(m86); daf-12(sa204*) double mutants (**Figure 6B**). FKH-8::GFP expression in the PDE dopaminergic ciliated sensory neuron is unaffected in *daf-19(m86); daf-12(sa204*) double mutants as assessed by co-localization with *dat-1:cherry (*91% PDE neurons are FKH-8::GFP positive in *wild type* animals and 92% in *daf-19(m86)* mutants, suggesting that FKH-8::GFP expression is unaffected in ciliated neurons (**Supplementary File 1**). *daf-19(m86)* allele affects all isoforms; as DAF-19 isoform D is expressed in ciliated neurons, our results suggest DAF-19D is not necessary for FKH-8 expression in ciliated neurons. In constrast, DAF-19 isoforms A and B seem to repress FKH-8 expression in non-ciliated neurons. *daf-19(of5),* a mutant allele that specifically affects *daf-19 a/b* isoform expression (**Figure 6A**) shows similar pan-neuronal de-repression of FKH-8::GFP (**Figure 6B**) further supporting the repressive role of DAF-19 A/B long isoforms.

Next, we aimed to assess if DAF-19 and FKH-8 act synergistically. *daf-19* and *fkh-8* genes are both located in chromosome II, despite several attempts, we failed to generate *daf-19(m86), fkh-8(vlc43) II; daf-12(sa204*) triple mutants but we were able to obtain *daf-19(m86), fkh-8(tm292); daf-12(sa204*) recombinant animals. We find DAF-19 and FKH-8 act synergistically in the regulation of *ift-20, peli-1, osm-1* and *xbx-1* reporters (**Figure 6C, D**). Of note, these reporters still show some vestigial expression in the triple mutant (**Figure 6D**). We CRISPR-engineered a full deletion of the *fkh-8* locus in the *daf-19(m86); daf-12(sa204); ift-20::rfp* strain which generated a viable triple null mutant [*fkh-8 (vlc39)* allele]. These animals show similar residual *ift-20* expression in a couple of unidentified neurons (**Figure 6C**), suggesting additional transcription factors might cooperate with DAF-19 and FKH-8 in the regulation of ciliome gene expression.

### Mouse FOXJ1 and FOXN4, master regulators of motile ciliome, can functionally replace FKH-8

Vertebrate FKH family is composed by 49 different members divided into 16 subfamilies (Shimeld et al., 2010). The establishment of specific orthology relationships between FKH members is challenging among distant species (Shimeld et al., 2010), precluding the direct assignment of the closest vertebrate ortholog for *C. elegans* FKH-8.

To date, no vertebrate FKH TF has been involved in ciliogenesis in primary cilia cell types. Nevertheless, in several vertebrate cell types that contain motile cilia, FoxJ1 FKH TF directly activates ciliome gene expression (Brody et al., 2000; Chen et al., 1998; Stubbs et al., 2008; Yu et al., 2008). Thus, considering its role in ciliogenesis, we next wondered if mouse FOXJ1 could functionally substitute FKH-8 in *C. elegans*. We find this to be the case as FoxJ1 cDNA expression under the dopaminergic promoter *dat-1* rescues *ift-20* expression similarly to *fkh-8* cDNA (**Figure 7A-C**). In *Xenopus*, another FKH TF, FoxN4, binds similar genomic regions to FoxJ1 and it is also required for direct ciliome gene expression in motile multiciliated cells (Campbell et al., 2016). We find FoxN4 expression also rescues *ift-20* expression defects in *fkh-8(vlc43)* animals. Importantly, we find that conserved functionality is not observed for any vertebrate FKH TFs as expression of mouse FoxI1, a FKH TF involved in the development of several tissues but not reported to control cilia gene expression (Edlund et al., 2015), does not rescue *fkh-8* mutant phenotype.

**Figure 7.**
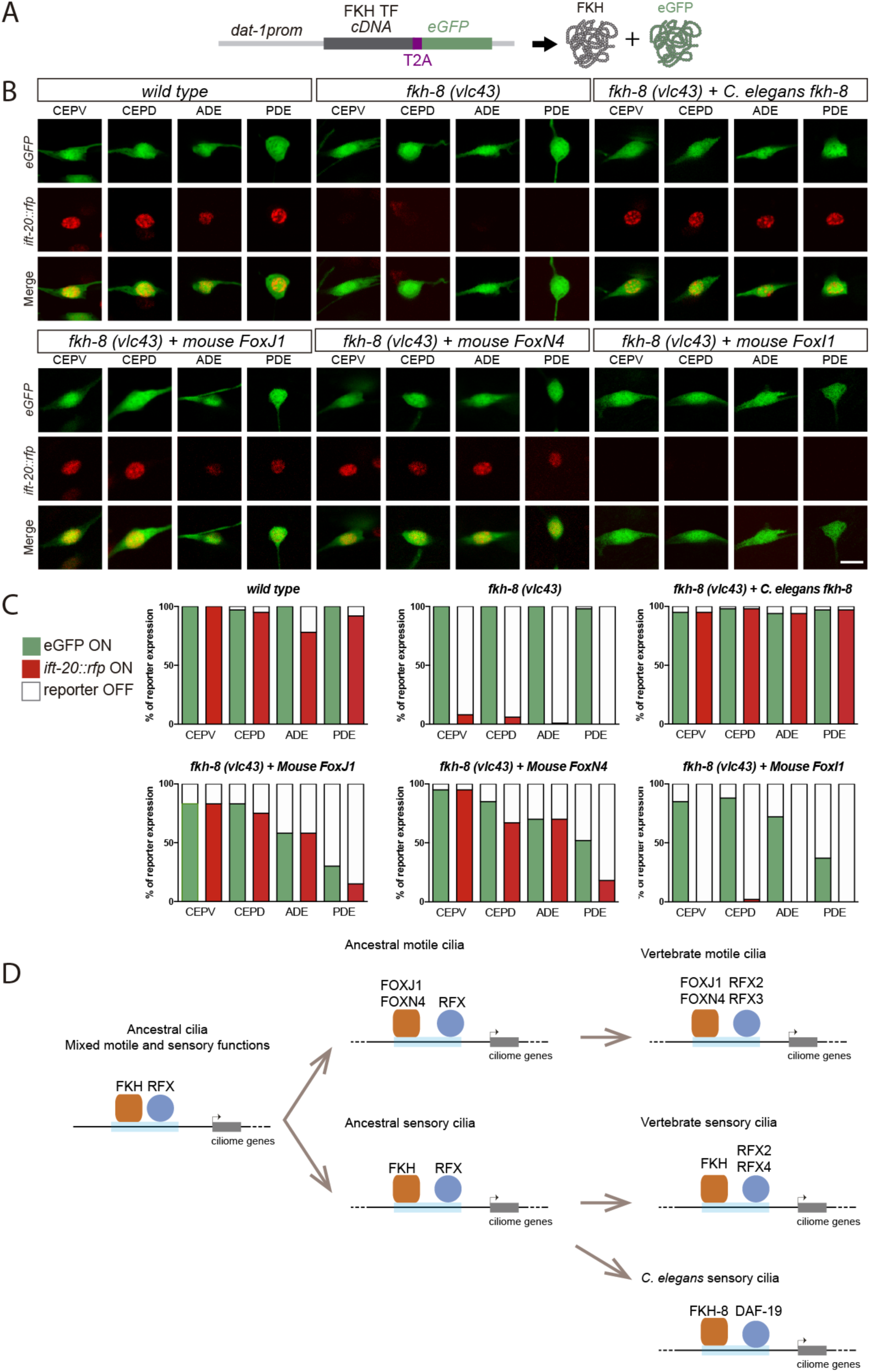
Mammalian FKH TFs with known motile cilia regulatory functions can rescue *fkh-8* mutant phenotype. A) Rescue strategy: *dat-1* promoter, unaffected in *fkh-8* mutants, is used to drive FKH TF cDNA and eGFP expression specifically in the dopaminergic ciliated system. B) Representative images of dopaminergic neurons expressing an integrated reporter for the core ciliome marker *ift-20* (in red) in *wild type*, *fkh-8(vlc43)* mutants and with the co-expression of different rescuing constructs. Scale bar = 5 μm. C) Quantification of rescue experiments. *ift-20* reporter expression is lost from the dopaminergic neurons in *fkh-8(vlc43)* null mutants compared to *wild type* animals. Expression of FKH-8, FOXJ1 and FOXN4 but not FOXI1 is sufficient to recover *ift-20* expression in dopaminergic neurons. N = 30 animals per transgenic line. See **Supplementary file 1** for similar results obtained with two additional transgenic lines per construct. D) Speculative model on the evolution of ciliome gene regulatory logic. FKH and RFX TFs could have an ancestral role in the direct coregulation of ciliome genes before its functional diversification into motile and primary cilia cell types. Different RFX and FKH TF members could have evolved to regulate ciliome genes in specific cell types in different organisms. Orange squares represent FKH TFs and blue circles RFX TFs, light blue bars represent ciliome enhancers.

In summary, our results unravel the functional conservation between FKH-8 and specific mouse members of the FKH family, which have already been described to work together with RFX TFs in the regulation of ciliome gene expression in motile cilia cell types.

## DISCUSSION

### FKH-8 works together with DAF-19 in the direct regulation of ciliome gene expression in sensory neurons

RFX are the only TFs known to be involved in the direct regulation of ciliome gene expression both in cell types with motile and sensory cilia. This role is conserved in nematodes, flies and vertebrates (Choksi et al., 2014). In this work we characterized the persistent activity of ciliome reporters in *daf-19/rfx* null mutants, demonstrating that, in some specific cellular contexts, DAF-19/RFX is not necessary to drive ciliome gene expression. DAF-19 is the only RFX TF in *C. elegans*; thus, persistent enhancer activity must be attributed to other TF families.

A multi-angled approach allowed us to identify FKH-8 as a key regulator of ciliome gene expression in most, if not all, sensory neurons in *C. elegans*. FHK-8 is expressed almost exclusively in all ciliated neurons and binds to upstream regions of many ciliome genes. *fkh-8* mutants show decreased levels of ciliome reporter gene expression, abnormal cilia morphology and defects in a plethora of behaviours mediated by sensory ciliated neurons. Finally, mutations in putative FKH binding sites for two ciliome reporters lead to expression defects, further supporting the direct action of FKH-8 in ciliome gene expression. Altogether, our results show that FKH-8 is a master regulator of ciliogenesis in sensory neurons and thus represents the first identified TF in any organism that works together with RFX in cell types with non-motile primary cilia.

In the past, the identification of direct targets of RFX TFs has been instrumental in the identification of new ciliome components, which lead to a better understanding of cilia function and the etiology of ciliopathies (Blacque et al., 2005; Chen et al., 2006; Efimenko et al., 2005; Li et al., 2004; Schiebinger et al., 2019). FKH-8 binds to many genes in the *C. elegans* genome, some with uncharacterized functions; thus, similar to RFX, a more exhaustive characterization of FKH-8 targets could be used to unravel novel components of the sensory ciliome.

### Specific DAF-19 isoforms repress *fkh-8* expression in non-ciliated neurons

Interestingly, our results show that DAF-19A and B isoforms repress (directly or indirectly) *fkh-8* expression in non-sensory neurons. Repression of alternative fates is a prevalent feature in neuronal development. DAF-19AB repression of *fkh-8* might be necessary to avoid ectopic ciliome gene expression in non-ciliated cells. Indeed, we find that, similar to *fkh-8*, *kap-1* gene, a core ciliome component involved in anterograde transport and cilia assembly is also pan-neuronally de-repressed in *daf-19* mutants (R.B. and N.F. unpublished). Repressive actions for DAF-19A/B have also been recently reported (Stasio et al., 2018).

### Role of FKH TFs in the transcriptional regulation of ciliome genes both in motile and sensory cilia cell types

Although TFs working with RFX in the regulation of ciliogenesis in sensory cell types where previously unknown, RFX TFs work in concert with the FKH TF FOXJ1 in the direct regulation of ciliome genes in different vertebrate cell types with motile cilia (Choksi et al., 2014).

Importantly, vertebrate sensory ciliogenesis is unaffected in FoxJ1 loss of function mutants (Choksi et al., 2014); thus, FoxJ1 role as a master regulator of ciliogenesis is restricted to motile ciliary cell types. In *Xenopus*, FoxN4 binds similar genomic regions to FoxJ1 and it is also required for motile ciliome gene expression (Campbell et al., 2016). We find both FOXJ1 and FOXN4, but not FOXI1, which has not been described to be involved in ciliogenesis, are able to functionally substitute FKH-8. This data suggests that specific FKH subfamilies might have an inherent capacity to act as direct ciliome regulators, independently of being expressed in motile or sensory cilia cell types.

### FKH-8 and DAF-19 show synergistic actions

FKH-8 bound regions are enriched for X-box/RFX sites, suggesting DAF-19 and FKH-8 are involved in the regulation of a common set of regulatory regions. Our double mutant analysis shows synergistic effects between *daf-19/rfx* and *fkh-8*, suggesting cooperative actions among these TFs.

Similarly, in motile multiciliated cells of *Xenopus* larval skin FOXJ1 binding to ciliome gene promoters depends on the presence of RFX2 (Quigley and Kintner, 2017). In addition, in human airway multiciliated epithelial cell, RFX3 and FOXJ1 act synergistically in the activation of ciliome genes (Didon et al., 2013).

Importantly, in *fkh-8, daf-19* double mutants, some enhancer activity is still present in several sensory cells underscoring the existence of additional direct regulators of ciliome gene expression. Nuclear hormone receptor *nhr-277* and *nhr-158* and the homeodomain *ceh-57,* whose expression is enriched in sensory neurons (**Figure 1F**), are possible candidates to be involved in the process and could be the focus of future studies.

### Evolution of cilia subtype specialization and ciliome regulatory logic

Ancestral cilium present in the last common eukaryotic ancestor has been proposed to combine motile and sensory functions (Mitchell, 2017). RFX role regulating ciliome expression predates the emergence of metazoans, where major cell type diversification has occurred (Chu et al., 2010; Piasecki et al., 2010). FoxJ and FoxN constitute the most ancient FKH sub-families, present in choanoflagelate *Monosiga brevicolis*, while FoxI subfamily is only present in bilaterians (Shimeld et al., 2010). Moreover, the ability of RFX and FKH TFs to bind similar genomic regions is not limited to metazoans and it is also present in fungi. In *Schizosaccharomyces pombe,* which lacks cilia and ciliome genes, Fkh2 FKH TF and Sak1 RFX TF bind the same regulatory regions to control cell cycle gene expression (Garg et al., 2015), suggesting that the joint actions for these TFs could be present before the split of fungi and metazoans. Alternatively, RFX and FKH TFs might have an inherent ability to cooperate that could explain convergent evolution of these TFs in ciliome regulation both in sensory and motile cilia cell types (Sorrells et al., 2018).

In light of these data, we hypothesize that RFX and FKH role as co-regulators of ciliome gene expression could precede the emergence of cilia division of labor and the specialization of motile and sensory cilium in different cell types (**Figure 7D**).

### Role of FKH TFs in ciliome regulation of primary cilia cell types

Regardless of the evolutionary history of events underlying RFX and FKH functions as master regulators of ciliome gene expression, our results raise the possibility that, in vertebrates, yet unidentified FKH TFs could work together with RFX as master regulators of ciliome gene expression in sensory ciliated cell types (**Figure 7D**). The establishment of specific orthology relationships between FKH members among distant species is challenging (Larroux et al., 2008; Shimeld et al., 2010) precluding the direct assignment of the closest vertebrate ortholog for *C. elegans* FKH-8. In addition, functional paralog substitutions among TFs of the same family have been described to occur in evolution (Tarashansky et al., 2021). Importantly, FoxJ1 and FoxN4 mutants do not show ciliome gene expression defects in non-motile ciliated cell types (Brody et al., 2000; Campbell et al., 2016; Chen et al., 1998; Stubbs et al., 2008; Yu et al., 2008). Other members of FoxJ and FoxN subfamilies are broadly expressed in mouse neurons, which all display primary cilia (Zeisel et al., 2018). It will be important, in future studies, to determine if additional FoxJ and FoxN TFs can rescue *fkh-8* expression defects in *C. elegans* and if they display similar roles in mammals as master regulators of sensory ciliome. These studies could also help better characterize the functional meaning of non-coding mutations associated to orphan ciliopathies.

## METHODS

### *C. elegans* strains and genetics

*C. elegans* culture and genetics were performed as previously described (Brenner, 1974). Strains used in this study are listed in **Supplementary File 4**.

### Mutant strain genotyping

Mutant strains used in this study are listed in **Supplementary File 4**. Deletion alleles were genotyped by PCR. Presence of *daf-19(m86)* allele was determined by visual inspection of the dye-filling defective phenotype of homozygous mutants. Presence of *daf-12(sa204)X* allele was ensured through a double cross strategy, crossing of F1 males with original *daf-12(sa204)X* mutants. Strains carrying point mutations were genotyped by sequencing. Genotyping primers are included in **Supplementary File 4**.

### DiD staining

Lipophilic dye filling assays were performed with the 1,1′-dioctadecyl-3,3,3′,3’ -tetramethylindodicarbocyanine, 4-chlorobenzenesulfonate salt (DiD) (Thermofisher, #D7757). DiD staining solution was freshly prepared prior to every assay as a 1:200 dilution of the DiD stock solution [2 mg/mL dilution in N,N-dimethyl formamide (Sigma, #D4551)] in M9 1X buffer. Animals were transferred into 1.5 mL tubes containing 200 μL of the DiD staining solution and incubated (wrapped in aluminium foil) for 2 hours at room temperature in an orbital shaker in a horizontal position. Animals were collected with a glass Pasteur and transferred to fresh NGM plates. Robust identification of the ASK, ADL, ASI, AWB, ASH, ASJ, PHA and PHB ciliated neurons was achieved through this method.

### Generation of *C. elegans* transgenic lines

Fluorescent reporters for ciliome genes were generated through fusion PCR (Hobert, 2002). To facilitate identification and scoring of reporter-expressing cells, GFP was tagged to the cell’s nucleus employing a modified sequence of the classical SV40 large T antigen nuclear localizing signal (NLS) (Kalderon et al., 1984). Regulatory sequences were amplified with custom oligonucleotides from N2 genomic DNA preparations. An independent PCR was used to amplify the 2xNLS::GFP::*unc-54* 3’UTR fragment from an NLS version of the pPD95·75 plasmid (pNF400). Successfully fused PCR products were purified using the QIAquick PCR Purification Kit (QIAGEN, #28106) and resuspended in nuclease-free water (Sigma, #W4502).

Mutated versions for the *xbx-1* and *ift-20* promoters were generated as PCR products by introducing the desired mutation of putative FKH sites within the corresponding custom primers. Putative FKH sites were identified through the single sequence scan tool from the CIS-BP website (Weirauch et al., 2014). Mutation criteria accounted for the nature of the nitrogenous bases and the number of hydrogen bonds they could form; thus, A was mutated to C and G was mutated to T (and vice versa). Mutated sequences were checked to discard the generation of new TF binding site motifs using both the motif scan tool of the CIS-BP database and the Tomtom tool (Gupta et al., 2007) from the MEME Suite website. When designed mutations created potential new TF binding sites manual punctual mutations were applied to disrupt those potential sites.

To generate FKH-8 rescuing plasmids, constructs containing the cDNA of the corresponding FKH TF fused to the self-cleaving peptide T2A (Ahier and Jarriault, 2014) and the eGFP cDNA from the pPD95.75 plasmid were created. Such constructs were then cloned under the control of the dopaminergic *dat-1* promoter between the KpnI/XhoI sites of the pPD95.75 backbone vector. *fkh-8* cDNA sequence was synthetically generated (Biomatik). Murine FKH members were obtained as Dharmacon clones (FoxJ1: MMM1013-202732974, FoxN4: MMM1013-211694291, FoxI1: MMM1013-202763055).

Simple-array transgenic lines were generated by intragonadal microinjection into strains of the appropriated genotype. The injection mix was composed by 50 ng/µL of a given purified fusion PCR or a rescuing plasmid plus 100 ng/µL of the pFR4 plasmid, *rol-6(su1006),* as a co-marker (Mello et al., 1991).

### Generation of *C. elegans* mutations

Whole deletion of the *fkh-8* locus was performed through a co-CRISPR strategy (Kim et al., 2014) using *dpy-10(cn64)* as conversion marker (Arribere et al., 2014). Custom CRISPR RNAs (crRNAs) were ordered (IDT, Alt-R® CRISPR-Cas9 crRNA XT) targeting both sides of the desired deletion of *fkh-8* and at the 5’ of the *dpy-10* site of mutation. Single stranded oligodeoxynucleotide (ssODNs) of approximately 100 base pairs overlapping each side of the genetic modifications were also ordered (IDT) and used as donor templates to achieve homology-directed repair. Cas9 nuclease (IDT, #1081058) and the universal trans-activating crRNA (tracrRNA) needed to initiate enzymatic activity (IDT, #1072532) were used. co-CRISPR injections were performed on young adult hermaphrodites expressing the reporter *otIs395(ift-20::NLS::tagRFP)III*. Microinjection mix was freshly prepared with all 3 crRNAs plus the tracrRNA, ssODNs and Cas9 nuclease. Ribonucleoprotein complex formation was achieved by incubating this mix for 10 minutes at 37 Celsius degrees. Before use, the final mix was incubated on ice for 30 minutes. All custom primer sequences and concentrations used for the generation of the aforementioned strains are included in the **Supplementary File 4**.

### Bioinformatical Analysis

Ciliome gene list was assembled including genes associated with cilium-related terms from the Gene Ontology using AmiGO (Carbon et al., 2009), known ciliome genes with functional X-boxes (Burghoorn et al. 2012) and genes whose expression in ciliated neurons was reported in the WormBase. Transcription factors were deliberately excluded from this list. A further curation process was performed through a bibliographic research (see **Supplementary File 2** for complete ciliome gene list).

For each isoform of the final 163 genes composing the ciliome gene list, putative regulatory sequences were retrieved from the Ensembl BioMart site (Kinsella et al., 2011) spanning 700 base pairs in length upstream of their translational start sites. These sequences were used to feed the RSAT oligo-analysis tool as previously described (Defrance et al., 2008; Turatsinze et al., 2008), using as a background model the in-tool genome of *C. elegans* and overall default options. Retrieved matrices were then compared both against the CIS-BP 1.02 (Weirauch et al., 2014) and the JASPAR core non-redundant 2018 (Khan et al., 2018) databases using the TomTom (Gupta et al., 2007) tool from the MEME suite (Bailey et al., 2009).

Identification of candidate transcription factors with enriched expression in ciliated neurons was performed through the on-line tool GExplore_1.4_ (Hutter and Suh, 2016), employing the sci-RNA-seq dataset by (Cao et al., 2017). A 5-fold enrichment ratio and a false detection rate of 0.001 were used.

Expression pattern data in each ciliated neuron type for candidate transcription factors at the fourth larval stage were retrieved from the *C. elegans* Neuronal Gene Expression Network (CeNGEN) (Taylor et al., 2021), whose results are freely accessible through the on-line tool SCeNGEA. Unfiltered data was used. ChIP-seq data from *C. elegans* TFs were retrieved from the ENCODE portal website (Davis et al., 2018) (time of consulting: January the 10th, 2019). Data analysis was performed through a custom script using R/Bioconductor (Huber et al., 2015). Peak annotation was carried out employing the ChIPseeker package (Yu et al., 2015), setting parameters as following: annotatePeak(gr1, tssRegion=c(-2000, 1000), level=lev, TxDb=annoData, overlap=“TSS”). ENCODE accession numbers for all datasets used in this analysis are listed in **Supplementary File 2**.

*fkh-8* ChIP-seq bed narrowPeak file (ENCODE accession: ENCFF653QKE) was used as input file for the web-based analysis tool ChIPseek (Chen et al., 2014). For de novo motif discovery, resulting fasta file with annotated peaks was then used to feed the RSAT peak-motifs tool as previously described (Thomas-Chollier et al., 2012b, 2012a), setting the number of motifs per algorithm at 10 and using all 4 available discovery algorithms with overall default options.

For gene onthology, genes associated to FKH-8 ChIP-seq peaks where analysed through the on-line tool WormEnricher (Kuleshov et al., 2016).

Gene expression correlation between TFs and genes of interest were calculated using embryonic sc-RNA-seq data (Packer et al., 2019). Genes of interest were categorized into four categories: 1) core ciliome genes, 2) subtype-specific ciliome genes (both extracted from our ciliome list), 3) panneuronal genes (Stefanakis et al., 2015) and 4) ubiquitously expressed genes (Packer et al., 2019). In addition to fkh-8 and daf-19, the proneural TF factor hlh-14 was added as control TF not related to ciliogenesis. For all 10,775 ciliated cells present in the dataset, correlation index (R) between the expression levels for each gene and the TF was calculated. R data for each gene category are represented in the graph (See **Supplementary File 2** for R values).

Presence of RFX/*daf-19* binding motifs within the FKH-8 ChIP-seq peak sequences was performed with the on-line tool Centrimo (Bailey and MacHanick, 2012) from the MEME suite. To prevent Centrimo from discarding sequences due to uneven sequence length within and among the different ChIP-seq datasets, a custom python script was used to extract sequences of 420 base pairs in length spanning 210 base pairs from the centre of each peak. This consensus length was used considering the average sequence length of FKH-8 ChIP-seq peaks. ENCODE accession numbers for all datasets used in this analysis are listed in **Supplementary File 2**.

Visualization and analysis of ChIP-seq and RNA-seq files were performed with the Integrative Genomics Viewer (IGV) software (Robinson et al., 2011).

### Microscopy

For scoring and image acquisition, worms were anesthetized with a drop of 0.5 M sodium azide (Sigma, #26628-22-8) on 4% agarose pads (diluted in distilled water) placed over a regular microscope glass slide (Rogo Sampaic, #11854782). These preparations were sealed with a 24 x 60 mm coverslip (RS France, #BPD025) and animals were then conveniently examined.

Scoring of ciliome features was performed observing the animals on a Zeiss Axioplan 2 microscope using a 63X objective. Assessment of fluorescence signal on PDE and Phasmid regions was performed *de visu*. To appropriately assess number of cells in the head, optical sections containing the volume of reporter-positive neurons in the head of the animals were acquired at 1 µm intervals and images were manually scored using FIJI (Schindelin et al., 2012). Reporters used in the FKH cis-mutational analyses (both *wild type* and mutated forms) were scored *de visu* as the low intensity and fast bleaching in their signals precluded us from taking pictures.

For cilia morphology assessment, image acquisition was performed with a TCS-SP8 Leica Microsystems confocal microscope using a 63X objective. The following conditions of optical sections (µm) were used: CEP: 0.4 µm; ADF: 0.2; AWB: 0.24; AWC: 0.3. Retrieved images were z-projected at maximum intensity (Leica LAS X LS) and linear adjustment for brightness and contrast was performed prior to cilia length quantification (N ≥ 32 cilia per neuron type) (FIJI). AWA analysis was performed from images acquired from dorsoventrally positioned animals (N = 7) in which both cilia were levelled and depth of arborisation was estimated from the volume containing all the optical sections (0.3 µm) in which fluorescence signal was observed.

All micrographs presented in this paper were acquired with a TCS-SP8 Leica Microsystems confocal microscope using a 63X objective and appropriate zooming conditions. Raw data and statistics for all scorings performed for this work are gathered in **Supplementary File 1**.

### Behavioural assays

Unless otherwise stated, all mechano- and chemosensory assays were performed over small-scale synchronized populations of young adult hermaphrodites.

Nose touch tests were performed as previously described (Kaplan and Horvitz, 1993). Ten minutes before the assay, young adult hermaphrodites were transferred to non-seeded NGM agar plates and nose touch responses were elicited by causing a nose-on collision placing an eyelash attached to a pipette tip in the path of an animal moving forward. With brief modifications from (Brockie et al., 2001), five consecutive nose touch trials were scored for each worm.

Both gentle and harsh touch mechanosensory tests were performed as previously described (Chalfie et al., 1985). Briefly, gentle touch assays were performed by alternatively stroking the animal just behind the pharynx and just before the anus with an eyebrow hair attached to a pipette tip for a total amount of 10 strokes (Hobert et al., 1999). Harsh touch assays were also performed by stroking the worms across the posterior half of their bodies in a top-down manner with a platinum wire. Each worm was tested five times with a 2 minutes interval between each trial (Li et al., 2011).

For all aforementioned mechanosensory assays, escape responses of trailed animals were recorded and a population response index (RI) was calculated for every replica as: RI = total number of escape responses / total amount of strokes Chemotaxis towards diacetyl, 2-heptanone, NaCl and 2-nonanone were performed over 3 times freshly washed worms with 1 mL of filtered, autoclaved CTX solution, aspirating the supernatant to a final volume of approximately 100 µL. 2 µL of this worm-containing solution with no less than 25 animals were placed at the proper place of the assay plates. During the assays, worms were allowed to freely crawl across the plates for 60 minutes at room temperature and then stored at 4 °C until the next day when worms’ positions were scored and behavioural indexes were calculated.

With few modifications, volatile diacetyl attraction assay was performed as described by (Margie et al., 2013). A four-quadrant paradigm drawn at the base of non-seeded NGM agar plates was used, adding a 1 cm circular central area that worms had to trespass to be scored. Stock diacetyl (Sigma-Aldrich, #803528) test solution was prepared as a 0.5% V/V mix in absolute ethanol (Scharlau, #ET00101000). Absolute ethanol was used as control solution. Immediately after the worms were plated, 2 µL of a mix combining equal volumes of diacetyl stock solution and sodium azide 1M were pipetted onto the 2 test sites (T) of the agar plate. Same procedure was then performed for the 2 control sites (C). Chemotaxis index (CI) was then calculated as: CI = (worms in (T1 + T2) - worms in (C1 + C2)) / total scored worms

Chemotaxis assay towards 2-heptanone was performed as previously reported (Zhang et al., 2016). A two-halves paradigm was used, adding the threshold distance by (Margie et al., 2013) to prevent immobile worms from skewing the data. 2-heptanone (Sigma Aldrich, #W254401) test solution was prepared as a 1:10 V/V mix in ethanol absolute. Ethanol was used as control solution. Immediately after the worms were plated, 3 µL of a mix combining equal volumes of 2-heptanone stock solution and sodium azide 1M were pipetted onto the test site (T) of the agar plate. Same procedure was follow to the control site (C). CI was calculated: CI = (worms in T - worms in C) / total scored worms.

Chemotaxis toward NaCl was also performed of a two-halves paradigm. Radial gradients of either test or control solutions were created prior to worm loading as originally stated (Ward 1973). Following (Frøkjær-Jensen et al., 2008), 10 µL of NaCl (Sigma, #S3014-1KG) 2.5 M (dissolved in double distilled water (ddH2O)) or ddH2O itself were respectively pipetted onto the agar surface at T and C spots and allowed to diffuse for 12-14 hours at room temperature. To increase steepness of the gradients, 4 µL of NaCl 2.5 M or ddH_2_O solutions were additionally added to the T and C spots respectively 4 hours prior to the chemotaxis assay. Chemotaxis indexes for two-halves paradigm assays were calculated as: CI = (worms in T - worms in C) / total scored worms.

Avoidance assay towards 2-nonanone was performed as previously reported (Troemel et al., 1997). Briefly, six equal sectors labelled as A, B, C, D, E and F were drawn on the base of squared plates (90 x 15 mm, Simport™, # 11690950) containing 15 mL of standard NGM agar. Stock 2-nonanone (Sigma-Aldrich, #W278550) test solution was prepared as a 1:10 V/V mix in absolute ethanol. Ethanol was used as control solution. Immediately after the worms were plated on the centre of the plate, 2 µL of a mix combining equal volumes of 2-nonanone stock solution and sodium azide 1M were pipetted onto two spots within peripheral test sector A. Same procedure was then performed for the ethanol control sites within sector opposite peripheral control sector F. Population avoidance index (AI) was calculated as: AI = (worms in (A+B) - worms in (E + F)) / total amount of worms.

Avoidance responses to water-soluble compounds were evaluated using the drop test as previously described (Hilliard et al., 2002). Following (Hilliard et al., 2004) with few modifications, well-fed synchronized young adult hermaphrodites were washed three times with M13 buffer. 5 animals were then placed on unseeded NGM agar plates and allowed to rest for 10 minutes. Two test solutions were assayed: 0.1% W/V sodium dodecyl sulfate (SDS) (Sigma, #L3771-100G) and 0.1 mM CuSO4 pentahydrate (Merck, #1027901000), both dissolved in the M13 buffer that acted as control solution. Each animal was tested first with 4 single drops of the control solution and then with 4 single drops of the testing solution, allowing for 2 minutes of recovery between each stimulus. Avoidance response was scored within 4 seconds after substance delivery. Population avoidance index (AI) per genotype and replica was calculated as: AI = number of responses / total amount of drops.

Dauer induction was performed using filtered liquid culture obtained from *wild type* worms grown at 7 worms/µl for 4 days. Briefly, 300µl of pheromone containing extracts or control extracts (culture media alone) were added to 60mm OP50-seeded NGM plates. After drying, 10 gravid worms were added and allowed to lay eggs for 18 hours and then removed from the plates. 72h later, resulting P0 worms were scored and percentage of dauer animals determined for each condition. Dauer induction was carried at 27°C in four independent experiments performed in parallel with wild type and *fkh-8(vlc43)* mutant worms.

Basal slowing response was performed with few modifications as previously reported (Sawin et al., 2000). In this case, 60 mm NGM plates in which HB101 was seeded in only one half of the plate were used. Briefly, well-feed worms were 3 times washed with 1 mL of filtered, autoclaved CTX solution, supernatant aspirated to a final volume of approximately 200 µL and 2 µL of this worm-containing solution (with no less than 10 animals) was placed at the non-seed part of pre-warmed assay plates. Free movement of the worms across the plates was recorded capturing 30 frames per second. Body beds per 20 seconds intervals were counted from same worms moving on agar and crawling across the bacterial lawn.

Sample size, tested genotypes, number of animals and number of replicates performed per assay are detailed in **Supplementary File 3**. All strains used for these behavioural studies are listed in **Supplementary File 4.**

### Statistical analyses

Statistical significance for the mean number of reporter-positive neurons in whole animals among different genetic backgrounds was assessed by the appropriate two-tailed t-test considering the homo- or heteroscedasticity of the samples being compared. Inbuilt Excel functions F.TEST and T.TEST were used and obtained p-values were adjusted through Bonferroni correction accounting for all possible pairwise comparisons in each experiment.

To increase for statistical power, statistical significance for the mean number of reporter-positive neurons in the five distinct anatomical regions containing ciliated neurons among different genetic backgrounds was assessed by the appropriate one-tailed t-test considering the homo- or heteroscedasticity of the compared samples. Obtained p-values were then adjusted through the Benjamini-Hochberg procedure setting α level at 0.05. This same procedure was used to assess for statistical significance within the dauer induction experiments.

Unless otherwise stated, same two-tailed t-test procedure was followed in the assessment of statistical significance in behavioural experiment. Behavioural responses were ultimately analysed through the corresponding indexes ranging from 0 to 1 (or to -1 to 0 when avoidance responses were assayed). For each type of assay, a population-based mean index was calculated per replica and a final response index was then obtained as the mean of all replicas’ means. Prior to hypothesis testing, the Shapiro-Wilk test (Shapiro and Martin, 1965) was used to address for the normality of these final indexes.

Assessment of synergistic effects between *fkh-8* and *daf-19* was performed under the multiplicative model (Wagner, 2015). Briefly, average number of reporter-expressing neurons found in the whole animals for each genetic background was transformed into the corresponding fold change related to the observed mean value in the *wild type*. Next, expected values for the fold change corresponding to triple *fkh-8*; *daf-12*; *daf-19* mutants were calculated as the product of the mean observed values for the double *daf-12*; *daf-19* and the single *fkh-8* mutant strains. Statistical significance between observed and expected values was then assessed through a one-sample t test.

For the assessment of statistical significance in rescue experiments, data was categorically classified as ‘on’ or ‘off’ and the significance of the association was examined using the two-tailed Fisher’s exact test. No further multiple testing correction was performed, as *fkh-8* null mutants were exclusively compared to wild type worms whereas each rescued line was exclusively compared against the *fkh-8* null mutants.

## ACKNOWLEDGMENTS

We thank CGC (P40 OD010440) for providing strains. Dr Laura Chirivella, Noemi Daroqui and Elia García for technical help. Erick Sousa for providing bioinformatical assistance. Ines Carrera and Elisa Martí for comments on the manuscript. Peter Swovoda for sharing *daf-19(of5)* allele.

## Funding

This work was supported by European Research Council (StG2011-281920 and COG-101002203), Ministerio de Ciencia e Innovación (SAF2017-84790-R and PID2020-115635RB-I00) and Generalitat Valenciana (PROMETEO/2018/055).

## AUTHOR CONTRIBUTIONS

R.B and N.F designed experiments, wrote the manuscript, and made figures with contributions from other authors. R.B conducted most of the experiments. A.E performed most behavioural assays, C.M and J.T helped in the bioinformatics analysis and C.M built fkh-8 CRISPR alleles.

## DECLARATION OF INTERESTS

The authors declare no competing interests.

---

**Supplementary Figure 1.**
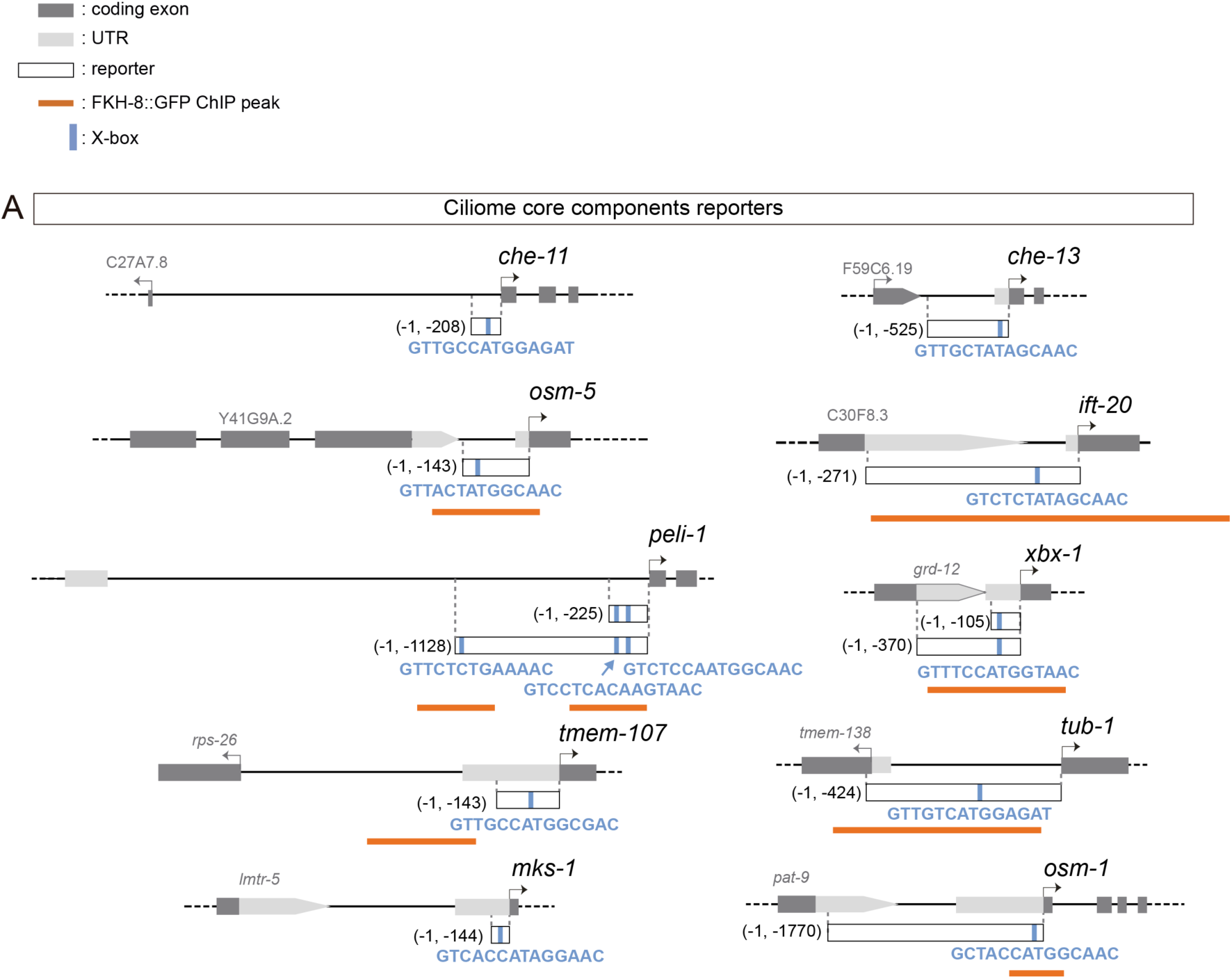
Ciliome reporters used in this work. Schematic representation of reporter constructs used in the manuscript. Selected core cilia components contain at least one experimentally validated X-box motif in their sequences (marked as a blue bar). For *che-11*, *che-13*, *osm5*, *ift-20*, *tub-1*, *mks-1* and *osm-1* see (Efimenko et al., 2005); for *peli-1* see (Chu et al., 2012), for *xbx-1* see (Schafer et al., 2003); for *tmem-107* see (Lambacher et al., 2016). Overlap between x-boxes and FKH-8 binding sites is found for *osm-5*, *ift-20*, *peli-1*, *xbx-1*, *tub1* and *osm-1*.

**Supplementary Figure 2.**
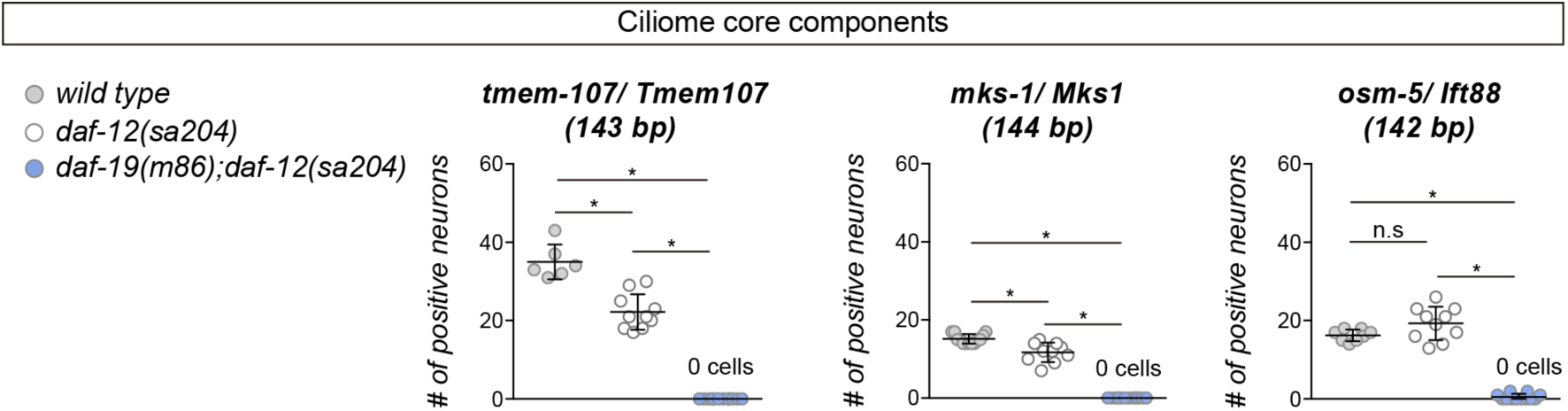
Lack of *daf-19* affects core ciliome expression. Expression of short reporters for the core cilia components *tmem-107*, *mks-1* and *osm-5* is completely abolished in double *daf-12(sa204)*; *daf-19(m86)* null mutants. *daf-12(sa204)* single mutants show slight but significant defects in *tmem-107* and *mks-1* reporter expression, and also for *che-13*, *ift-20* and *osm-1* (not shown in the graph). Each dot represents the total number of reporter-positive neurons scored in a single animal. Mean and standard deviation are represented. See **Supplementary file 1** for raw scoring data in all genetic backgrounds and for all reporters.

**Supplementary Figure 3.**
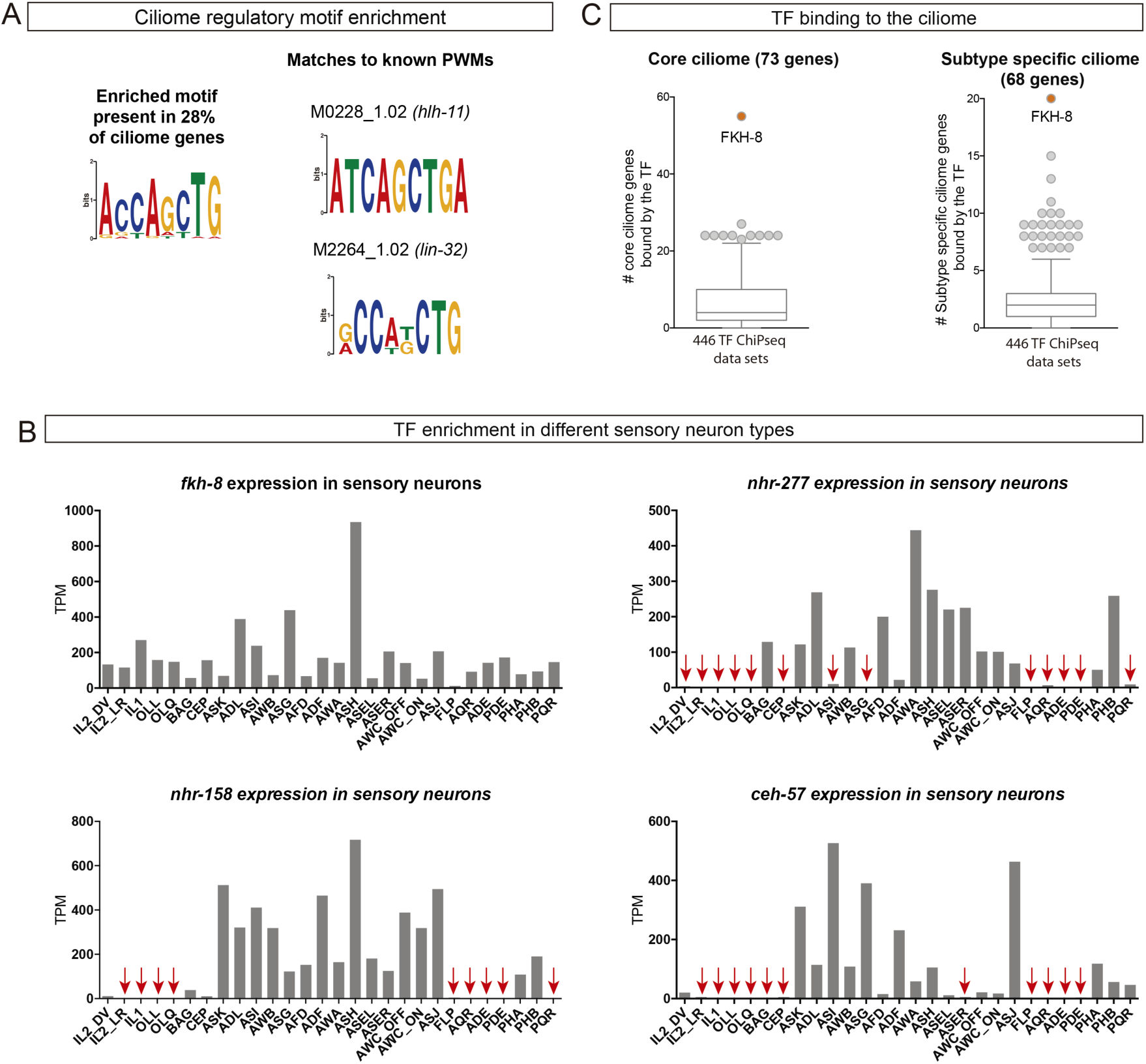
Available -omics data identifies FKH-8 as a candidate transcriptional regulator of ciliome genes in *C. elegans*. A) *De novo* motif enrichment analysis of putative regulatory sequences of ciliome genes identifies a motif matching known binding site for the bHLH TFs *lin-32* and *hlh-11*. B) sc-RNA-seq data of FACS-isolated neurons from L4 hermaphrodites (Taylor et al., 2021) show broad expression for *ceh-57*, *fkh-8*, *nhr-158* and *nhr-277* TFs across the whole ciliated system of *C. elegans*. Only *fkh-8* expression is detected in all ciliated neuron types. Red arrows indicate values lower than 10 TPM (transcripts per million). C) ChIP-seq data analysis shows FKH-8 ranks first among 259 TFs directly binding to either core ciliome genes (left) or subtype-specific ciliary features (right).

**Supplementary Figure 4.**
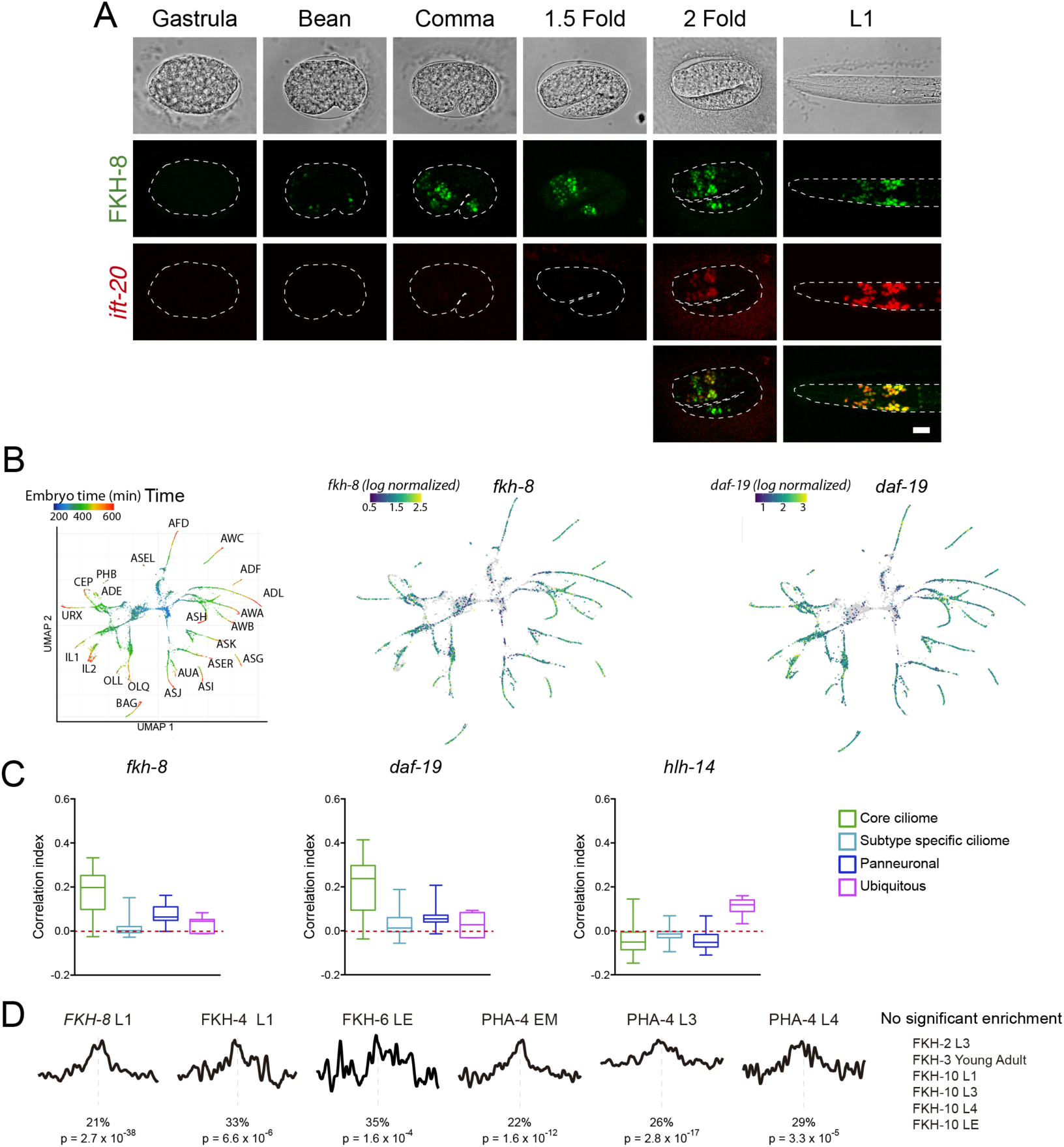
*fkh-8* expression along development of the ciliated system. A) Representative images of developmental embryonic milestones until hatching (L1) in animals expressing both a fosmid-based reporter for *fkh-8* (in green) and an integrated reporter for the panciliary marker *ift-20* (in red). Note that due to long maturation time of the tag-RFP reporter, *ift-20::tagRFP* expression is only detected from the 2 fold stage, while *ift-20::gfp* reporter in Figure 2 is first detected at bean stage, similar to *fkh-8* expression. Scale bar = 10 µm. B) Embryonic sc-RNA-seq data (Packer et al., 2019) from *C. elegans* ciliated neurons. Pseudo-time (left pannel) shows the maturation trajectory of ciliated neurons that coincides with increasing *fkh-8* (centre) and *daf-19* (right) expression. C) Correlation index between *fkh-8, daf-19* and *hlh-14* TF scRNAseq expression and genes divided in four different categories (core ciliome, subtype ciliome, panneuronal or ubiquitous) for all the ciliated lineages (Packer et al., 2019). *fkh-8* and *daf-19* expression shows high correlation index with core ciliome genes but not for other gene categories, while *hlh-14*, bHLH TF not involved in ciliogenesis shows low correlation values in all categories. See **Supplementary file 2** for raw data. D) Presence of DAF-19/RFX binding motifs is less significantly or not significantly enriched in ChIP-seq datasets for other FKH TFs. See **Supplementary file 2** for detailed data.

**Supplementary Figure 5.**
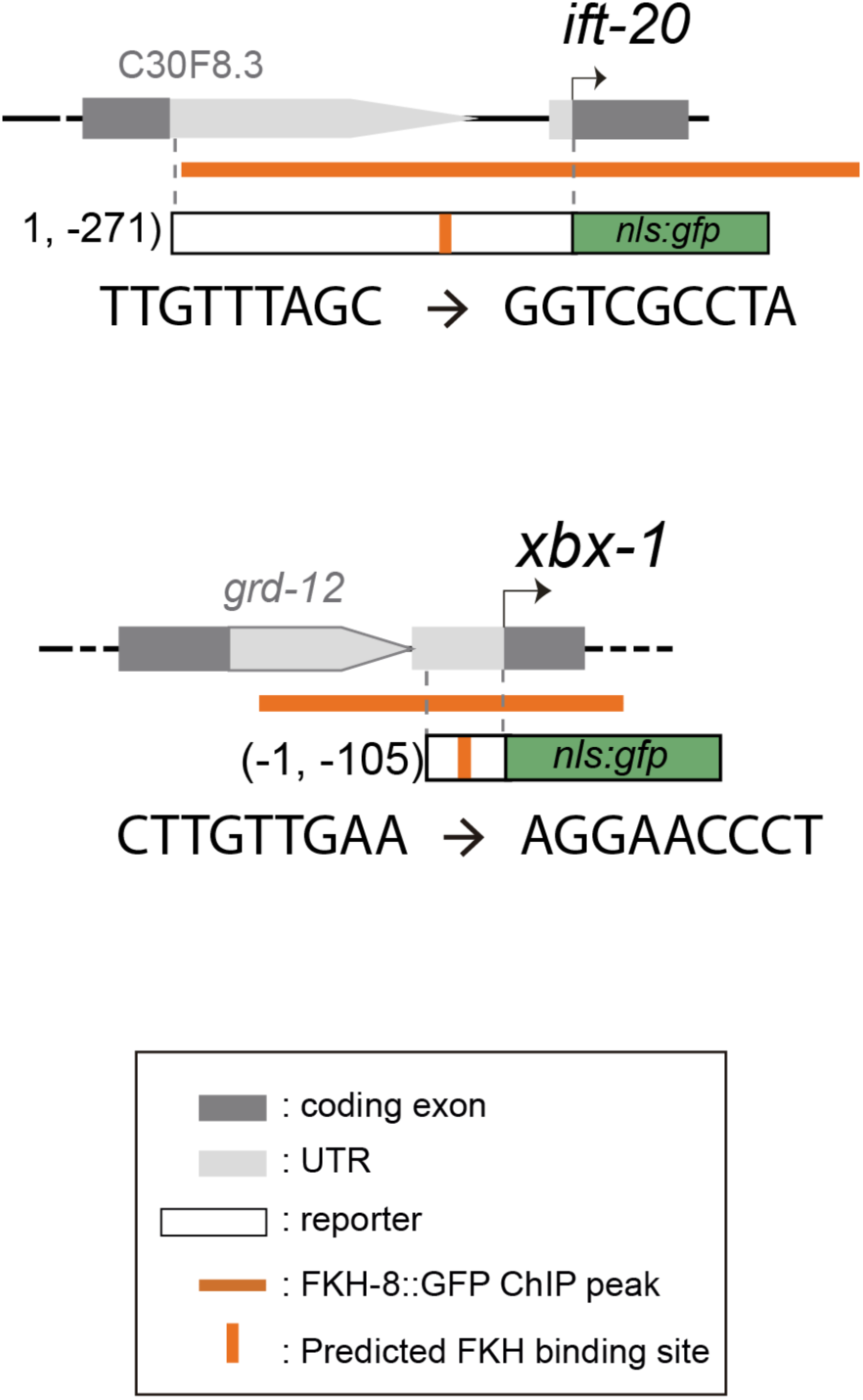
*cis*-mutation of putative FKH sites of two core ciliome components. Schematics for the *ift-20* and *xbx-1* loci and reporters. Dark grey boxes represent exons whereas light grey boxes correspond to UTRs. FKH-8 peaks are depicted with an orange horizontal line while predicted FKH DNA binding motifs are indicated with a vertical orange bar. Sequences corresponding to *wild type* and mutated putative FKH sites are indicated. See **Supplementary file 1** for raw scoring data and **Supplementary file 2** for FKH putative binding site assignment.

**Supplementary Figure 6.**
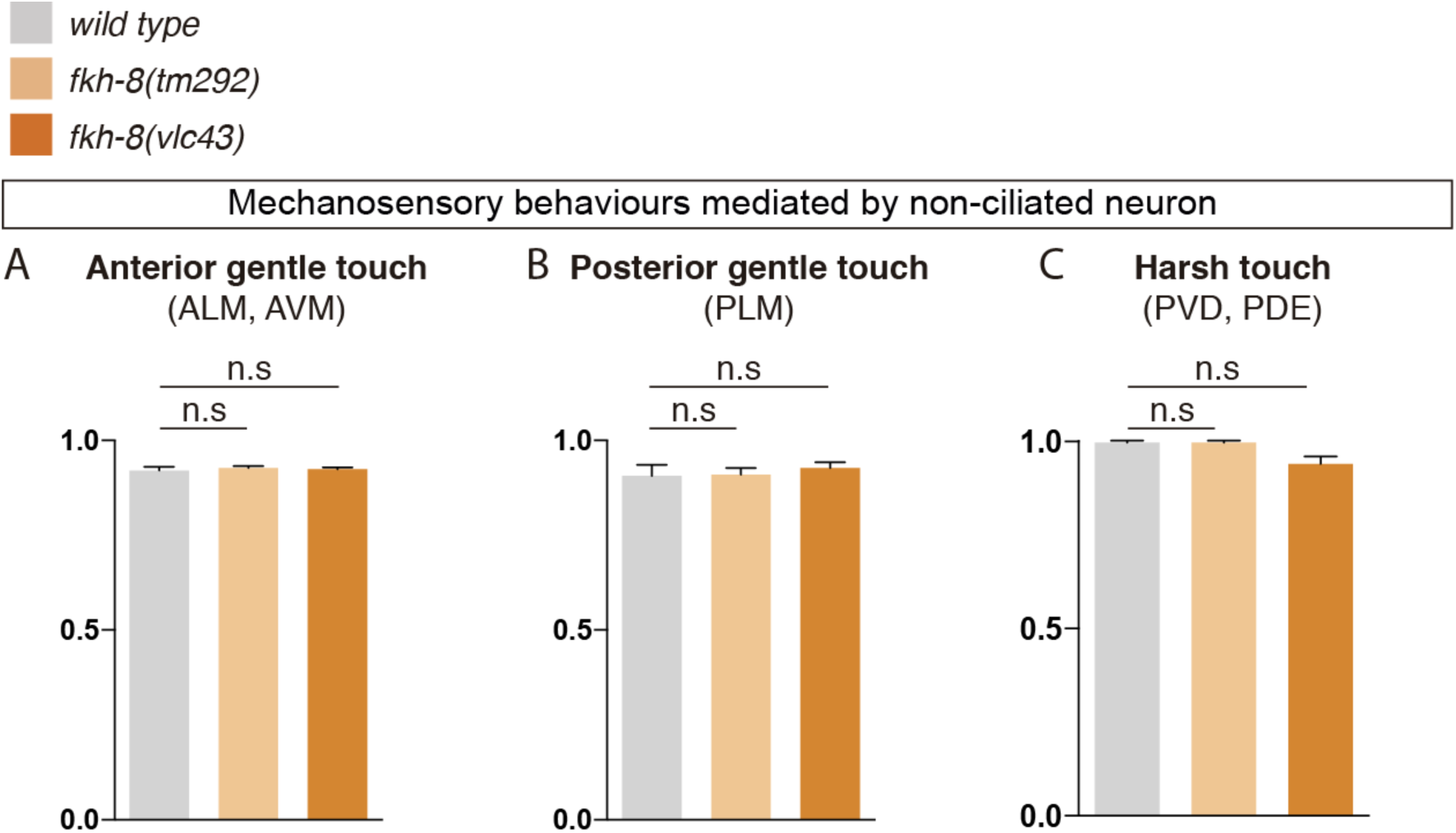
FKH-8 is not required for correct display of mechanosensory behaviours mediated by non-ciliated neurons. A to C) *fkh-8* mutants show normal avoidance behaviours elicited by mechanical stimuli known as gentle touch and harsh touch paradigms, suggesting FKH-8 is not required for the correct functionality of non-ciliated neurons ALM, AVM, PLM and PVD. Redundant actions of PVD and PDE controlling scape response to harsh touch prevent to assess defects about the functionality of ciliated PDE neurons. Mean and standard deviation are represented. See **Supplementary file 3** for raw data and samples’ sizes.

**Supplementary Figure 7.**
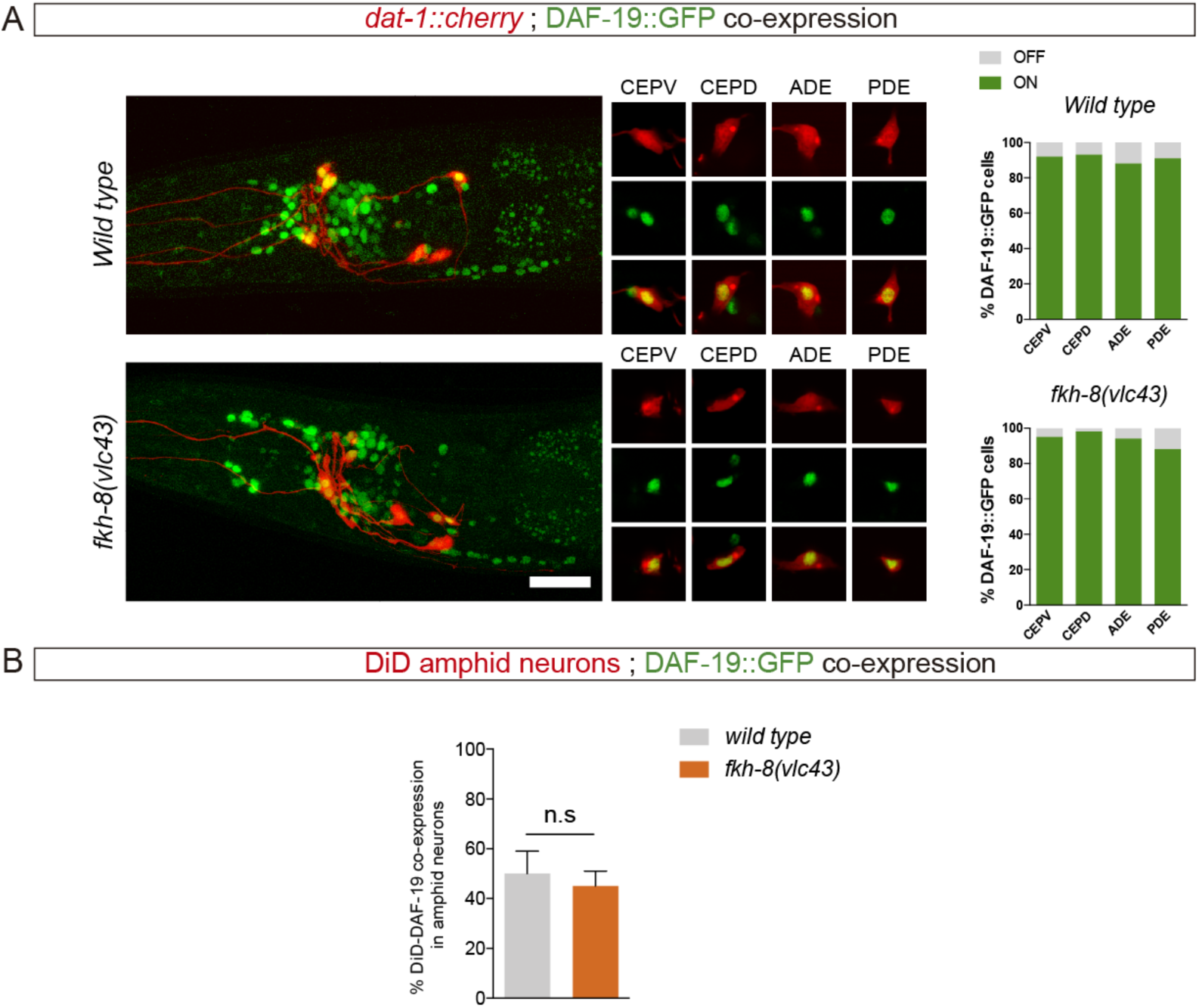
Lack of FKH-8 has no major effect on DAF-19 expression. A) Representative lateral views from heads of young adult hermaphrodites co-expressing a fosmid-based DAF-19::GFP reporter and *dat-1::mcherry* reporter labelling the dopaminergic neurons. Lack of FKH-8 does not seem to affect DAF-19::GFP expression pattern. Co-localization analysis shows normal expression in the dopaminergic ciliated neurons (CEPV, CEPD, ADE, PDE), quantified in the graphs. Scale bar = 20 µm. See **Supplementary file 1** for raw data and samples’ sizes. B) *daf-19* expression is largely unaffected in the subpopulation of DiD-positive ciliated amphid neurons in null *fkh-8* mutant animals. DAF-19::GFP is consistently detected in the ASI, ADL and AWB neurons in both *wild type* and null *fkh-8* mutant backgrounds. Mean and standard deviation are represented. N = 10 animals.

## Notes

### Competing Interest Statement

The authors have declared no competing interest.

